# Allopatric montane wren-babblers exhibit similar song notes but divergent vocal sequences

**DOI:** 10.1101/2022.06.25.497582

**Authors:** M. Abhinava Jagan, Ananda Shikhara Bhat, Anand Krishnan

## Abstract

The songs of passerine birds consist of notes temporally arranged into vocal sequences following syntactic structures, and function both in courtship and territorial defense. Geographic barriers are important drivers of avian speciation, and also influence the divergence of song. However, there is relatively little quantitative study of the relationship between geographic barriers and the syntactic structure of vocal sequences. Here, we investigate interspecific divergence in song notes and syntax within a genus of allopatric montane Asian wren-babblers (*Spelaeornis*). Employing a robust quantitative analytical framework and song recordings from publicly accessible databases, we find that *Spelaeornis* appear to have undergone diversification in song syntax without divergence in note parameters. Broadly, we find three different syntactic structures across the eight species in the genus, each occurring in a different geographic region in Asia, with two species apparently exhibiting intermediate syntax. Species within the genus appear to possess similar song notes, but subgroups confined to different geographic regions (eg: hills south of the Brahmaputra river) arrange these notes according to different syntactic rules to construct songs. Our computational framework to examine signal structure and diversification across multiple scales of signal organization has potential implications for our understanding of speciation, signal evolution and more broadly in fields such as linguistic diversification.

## Introduction

Allopatric speciation, the process underlying the formation of new species by geographic isolation, has long been a subject of research interest (Price 2007). Geographic barriers such as rivers, mountains, oceans or valleys can isolate populations of a species, thereby preventing gene flow. This reproductive isolation, over time, results in accumulation of genetic differences, such that the two species remain isolated from each other on secondary contact (Irwin et al. 2001). The initial reproductive isolation may be reinforced by pre-mating isolation mechanisms such as temporal separation of breeding periods (Hillis 1981; Marshall and Cooley 2000), divergent habitats (ecological isolation) (Egan and Funk 2009), morphological differences (mechanical isolation) (Richmond et al. 2011) and behavioral differences (behavioral isolation) (Marler and Slabbekoorn 2004; Uy et al. 2018). Of these, the last is particularly relevant to organisms that learn behavioral traits, because these learned traits may diverge relatively rapidly over evolutionary time either for adaptive reasons or through neutral evolutionary processes (Lachlan and Servedio 2004; Yeh and Servedio 2015). Understanding allopatric speciation thus requires a comprehensive understanding of how behaviors and sensory signals vary among close relatives and across geographic barriers.

Perhaps the best-studied learned signaling traits in this regard are the complex, culturally transmitted songs of passerine birds. This learned behavior exhibits complex structures that, among other communicative functions, support species recognition and minimize interspecific hybridization (Marler and Slabbekoorn 2004; Bradbury and Vehrencamp 2011). Each song consists of individual vocal units - notes or syllables, which are ordered in complex temporal arrangements to form vocal sequences. This temporal arrangement is referred to as the syntax of the song (Marler and Peters 1988; Kershenbaum et al. 2012; Fishbein et al. 2020). Syntax may follow a number of rules, including repetitive songs, repetitions of specific motifs, or highly complex songs consisting of a number of note types. Both the acoustic structure of individual notes and the syntactic structure of vocal sequences play an important role in mate attraction, species recognition, and communication of context in diverse vertebrate taxa (Peters et al. 1980; Searcy et al. 1981; Searcy and Marler 1981; Balaban 1988; Marler and Peters 1988; Isler et al. 1998; Nowicki et al. 2001; Wanker and Fischer 2001; Charrier and Sturdy 2005; Dahlin and Wright 2012; Kershenbaum et al. 2012; Briefer et al. 2013; Kershenbaum et al. 2016; Suzuki et al. 2018; Ciaburri and Williams 2019; Engesser and Townsend 2019; Leroux et al. 2021; Bhat et al. 2022). Thus, it is important to study acoustic signals at multiple hierarchical scales of organization to understand their evolution and function.

A number of studies have shown that geographic barriers exert potent influences on the structure of bird song (Marler and Tamura 1962; Baker 2006; Baker et al. 2006; Podos and Warren 2007; Irwin et al. 2008; Kirschel et al. 2009; Grant and Grant 2010; Lachlan et al. 2013; Lachlan et al. 2016). In white-crowned sparrows, geographically separated populations exhibit different dialects (Marler and Tamura 1962), and the same pattern is observed in insular populations of birds separated from the mainland (Baker et al. 2006; Lachlan et al. 2013). Song structure could evolve by simple addition and deletion of new notes to the repertoire, through changes in the syntactic rules which dictate how songs are constructed (i.e. increased repetition of the same notes, use of certain motifs, etc.), or through a combination of both these processes (Fig 1A). Quantitative studies that examine both notes and syntactic structure of song in allopatric bird species are therefore needed to better understand the role of note change versus sequence change in the evolution of song. Such an approach would enable us to examine how syntax changes across geographic barriers in comparison to divergence in note structure. For example, do geographically proximate species exhibit similar syntactic rules but different notes, or different syntactic structures with or without note change? Given the parallels between bird song and human language (Hunt 1923; Doupe and Kuhl 1999; Salwiczek and Wickler 2004; Berwick et al. 2011; Miyagawa et al. 2013; Collier et al. 2014; Henry et al. 2015; Sainburg et al. 2019), such research also helps understand how geography potentially influences human linguistic evolution.

**Fig 1:**
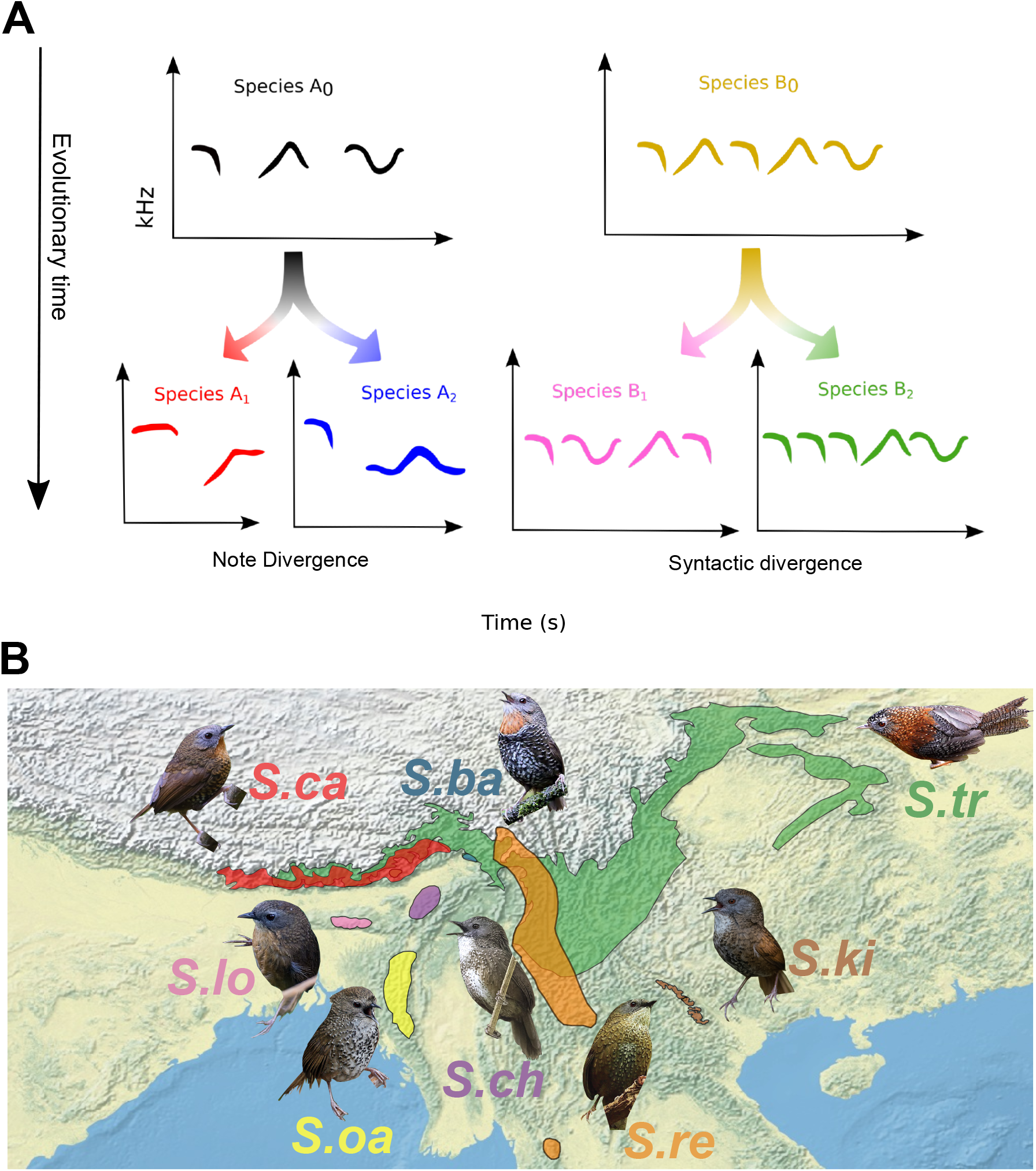
**A**. Acoustic signal divergence may involve divergence in the frequency-time parameters (acoustic structure) of individual notes (note divergence, left), and the rules followed to order these individual notes into a song (syntactic divergence, right). **B**. Geographical ranges of the genus *Spelaeornis*, and the abbreviated names we use throughout. *S*.*ca: S. caudatus, S. ba: S. badeigularis, S. tr: S. troglodytoides, S. ch: S chocolatinus, S. re: S. reptatus, S. ki: S. kineari, S. oa: S. oatesi, S. lo: S. longicaudatus*. The species are allopatric, and *S. tr* is present at higher altitudes than its congeners. The images of *Spelaeornis* were sourced with permission from the Macaulay library (Cornell Laboratory of Ornithology, Ithaca, NY, USA) (ML101191261- Jon Irvine, ML379201951- Anupam Nahardeka, ML379618361-Lakpa Tenzing Sherpa, ML378245821- Pranjal J. Saikia, ML213328961- Xiwen Chen, ML269988921- Ngoc Sam Thuong Dang, ML205760231- James Eaton, ML378358871- Firoz Hussain). Species ranges were sourced from International Union for Conservation of Nature (IUCN 2019) and the figure was created in QGIS (Quantum GIS Development Team 2013; http://qgis.osgeo.org).

*Spelaeornis* is a genus of montane understory birds from Asia containing eight species with non-overlapping ranges (del Hoyo et al. 2014). All species of this genus are endemic to montane forest from the Himalayas to South-East Asia (see Methods, Fig 1B) (IUCN 2019). They thus present a good system to comparatively examine how the geographic barriers between hill ranges have shaped the evolution of vocal sequences. Here, we quantitatively examine the syntactic structure of vocal sequences as well as the parameters of song notes in this genus. Specifically, we first ask whether individual song notes vary between species. Secondly, we quantitatively analyze the syntactic structure of *Spelaeornis* songs to address whether different allopatric species exhibit distinct vocal sequences, arranged according to different rules. We also hypothesized that that geographically proximate species possess similarities in their syntactic structure and vocal sequence content. Our study thus examines how geographic barriers influence diversification in complex learned signals across multiple organizational scales.

## Materials and Methods

### Study species

The genus *Spelaeornis* consists of eight species of allopatric montane wren-babblers- *S. caudatus* (*S. ca*) endemic to the Eastern Himalayas, *S. badeigularis* (*S. ba*) of the Mishmi Hills in India, *S. troglodytoides* (*S. tr*), that replaces other *Spelaeornis* species at higher elevations, *S. chocolatinus* (*S. ch*), *S. oatesi* (*S. oa*), and *S. longicaudatus* (*S. lo*) of hill ranges south of the Brahmaputra river, *S. kinneari* (*S. ki*) and *S. reptatus* (*S. re*) from Southeast Asia (Fig 1B) (Rasmussen and Anderton 2005; Collar 2006; King and Donahue 2006; del Hoyo et al. 2014). These allopatric species are endemic to restricted ranges, replacing each other in wet montane forests. They therefore provide us with a good model system to examine the effect of geographic barriers on interspecific divergence in acoustic signals.

### Recordings

We sourced song recordings of the genus *Spelaeornis* from multiple online song databases including Macaulay library (https://www.macaulaylibrary.org/), Xeno-Canto (https://xeno-canto.org/), and AVoCet (https://avocet.integrativebiology.natsci.msu.edu/) (see Supplementary Data 1 for a full list of recordings, including recordist and location). This dataset was curated to include recordings that were longer than 10 seconds in duration with relatively low background noise, and also to ensure that none of the recordings were duplicates from different databases. For purposes of sample size, we considered a ‘recording’ as being all audio database files recorded by the same recordist on the same day, and also the total number of songs measured per species and the number of notes. Our total sample for analysis was thus as follows: *S. ca*: 1864 notes, 244 songs analyzed from 28 recordings; *S. ba*: 1358 notes, 253 songs from 14 recordings; *S. tr*: 1574 notes, 213 songs from 19 recordings; *S. ch:* 759 notes, 135 songs from 12 recordings; *S. re*: 622 notes, 50 songs from 4 recordings; *S. ki:* 904 notes, 62 songs from 8 recordings; *S. oa:* 1561 notes, 219 songs from 14 recordings; *S. lo:* 940 notes, 156 songs from 11 recordings.

### Comparison of note parameters

Using Raven Pro Version 1.6 (Cornell Laboratory of Ornithology, Ithaca, NY, USA), we first digitized individual notes of each species and calculated 10 parameters that described the properties of each note: note duration, 90% bandwidth (difference between the frequencies containing 90% of energy in the power spectrum), peak frequency, minimum and maximum frequencies of the peak frequency contour, start and end frequencies of the peak frequency contour (these combined giving parameters of note shape), center frequency (frequency of 50% energy in the power spectrum), average entropy, and the time of maximum energy relative to the start of the note (peak time relative). We then performed a Principal Components Analysis (PCA) on the correlation matrix of these parameters, to examine interspecific note overlap of *Spelaeornis* in signal space. To quantify this overlap, we trained a Linear Discriminant classifier on the note parameters described above, using the Classification Learner app in MATLAB (Mathworks Inc., Natick, MA, USA) with 10-fold cross-validation, and assessed the accuracy with which notes were correctly assigned to the different species. We additionally performed a statistical randomization test of whether species overlapped more or less than expected by chance. For this, we randomly reshuffled data points across species, maintaining the sample size for each individual species (Chek et al. 2003; Luther 2009; Schmidt et al. 2013; Krishnan 2019; Chitnis et al. 2020). For each of 1000 such randomized datasets, we calculated the average interspecific Euclidean distance (for the first three principal components) in signal space. Finally, we compared the observed average interspecific distance to the distribution obtained from the randomized datasets by computing a *Z*-score. If the song notes of *Spelaeornis* overlapped in signal space, the observed interspecific distance in the signal space should be less than that expected by chance alone, thus resulting in a negative *Z-*score.

### Classifying note types and note groups

Before analyses of song syntactic structure, we constructed sequences illustrating the temporal structure of each song. This required us to first classify song note types for each species based on differences in spectral structure and the parameters we measured (Fig 2A). We verified these note type classifications in two ways: first, by measuring classification accuracy using a linear discriminant classifier as detailed above, and secondly, by cross-verifying the note classifications using a second observer in a double-blind paradigm. In a small number of cases, the classifier incorrectly assigned visually dissimilar notes to the same group, and we manually defined the notes as subclasses of the same note type in these cases. However, this was rare (three instances), and the combination of methods generally enabled us to reliably classify note types. We used these note types to calculate measures of song complexity (see below). However, note type classifications did not permit interspecific comparisons of syntax as the note types and diversity were not statistically comparable across species (subsequent analyses required that all species be compared within the same state space). We therefore also grouped notes into “note groups’’, which were consistent across species, based on their spectral shape (Fig 2A). This enabled direct interspecific comparison of song syntax, as notes with minor variations in spectral structure or frequency were pooled into the same note group. As before, we verified note group classifications in a double-blind paradigm (i.e., a second author also scored note groups). Across species, we thus hierarchically grouped notes into nine note groups, abbreviated as letters in the relevant figures and tables. These were: Ascending (a), Descending (b), Ascending-Descending (c), Descending-Ascending (d), Ascending-Descending-Ascending (e), Descending-Ascending-Descending (f), Notes with constant frequency (k), Complex notes with more than two inflection points and a total duration of less than 200 ms (g), Long complex notes with more than two inflection points and a total duration of more than 200 ms (l). To ensure that we had adequately sampled all the note types/note groups, we computed note accumulation curves for each species by randomizing the order of digitized notes for all songs.

**Fig 2:**
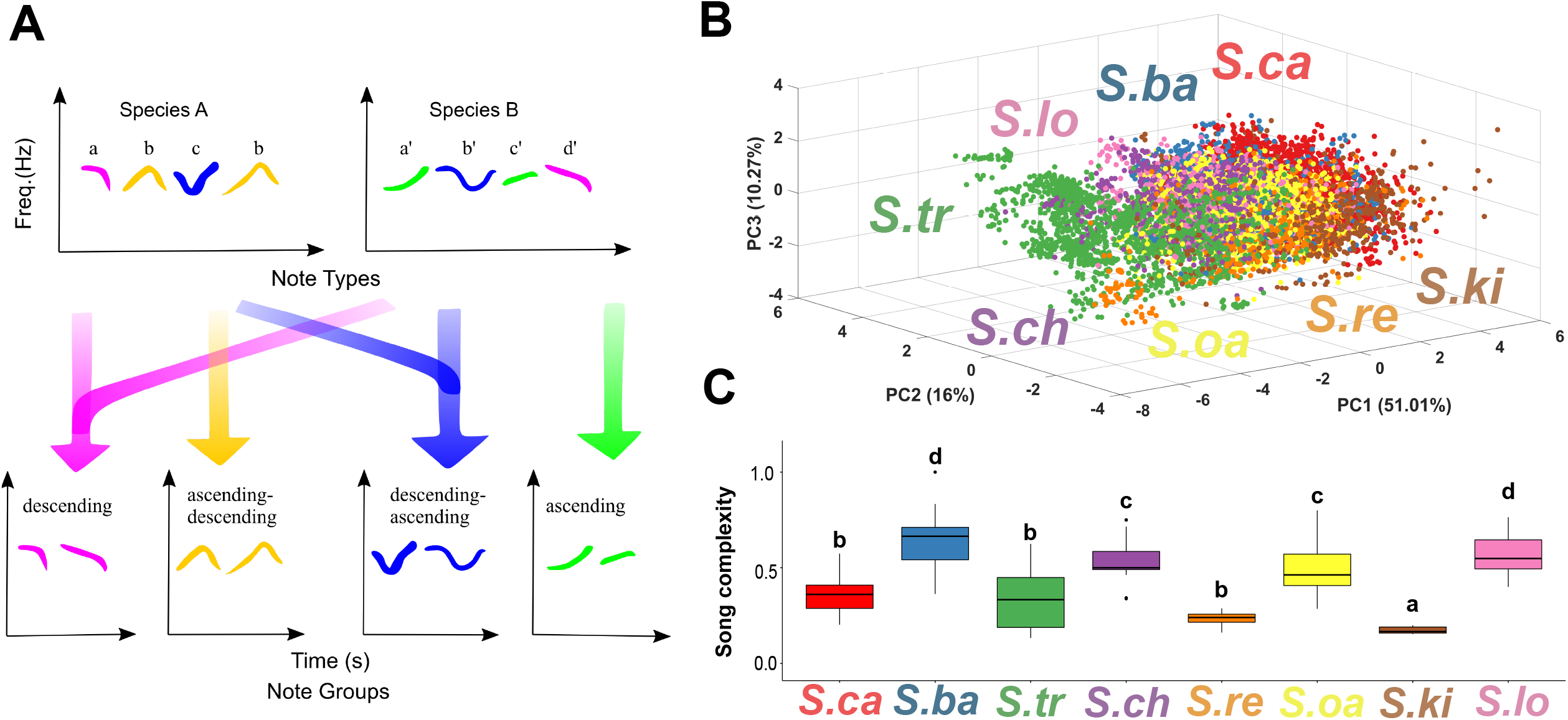
**A**. Schematic representation of the framework followed to classify notes from different species into note types and note groups (see Methods). Different note types of Species A and Species B are depicted by alphabets (top), and different note groups in each species are represented by different colors. Four examples of note groups present in Species A and Species B are described in the bottom panel. The Y-axis represents the frequency of each note, and the X-axis represents time. **B**. Three-dimensional signal space of *Spelaeornis* constructed using the first three principal components (PC 1-3) of the correlation matrix of 10 measured note parameters. Each point in this space represents a note, and the species are represented by different colors. *Spelaeornis* exhibit overlap in note parameter space. **C**. Box plot depicting the song complexity metric across species in *Spelaeornis*. Letters depicted represent groups showing statistically significant differences after pairwise comparisons. a: differed significantly from *S. oa, S. lo, S. ch* and *S. ba*; b: differed significantly from *S. ba* and *S. lo* but not from *S. oa, S. ch, S. ki*; c: differed significantly only from *S. ki*; d: differed significantly from *S. ca, S. tr, S. re and S. ki*. Further details are given in the Results and Table S2.

### Syntactic analyses

To examine and compare vocal sequences and syntactic structures, we employed a series of metrics calculated using both note types and note groups. Using the note type classifications, we first calculated a metric of song complexity- the number of different note types in a song divided by the total number of notes in the same song. For each song (sample sizes above) for each species, we computed and compared the average values of song complexity for each recording (the unit “recording” defined above) using a Kruskal-Wallis test with a post-hoc Dunn test. Because further syntactic comparisons relied on species having similar sample spaces, we used note groups as a unit in subsequent analyses for direct interspecific comparison (although we also repeated these analyses using note types to ensure that the type of classification did not influence the patterns observed). To examine song syntax in *Spelaeornis* we constructed note group sequences for each song, for each species. We first modeled *Spelaeornis* song structure as a first order Markov chain and constructed a transition probability matrix (*N*) for each species, where the i-j^th^ entry of the matrix, *N*_*ij*_, represents the probability of transition from note group i to note group j. To further validate our analysis, we employed a co-occurrence metric which is free of Markovian assumptions (Bhat et al. 2022). This analysis is based on the premise that not all vocal sequences in animals may be Markovian (Kershenbaum et al. 2014), and it is thus important to cross-verify results without making underlying assumptions about the processes involved. Here, for a given set of note group sequences, we define ^d^C_ij_ as the probability that the note group j occurs within d-1 notes of note group i. We computed ^d^C_ij_ for song sequences for each species. Next, we constructed artificial sequences where notes were randomly distributed following a stationary distribution. In this stationary distribution, the probability of a note occurring in a sequence was given by the proportion of occurrence of that note in the species’ song. We generated 50 such artificial sequences to compute a robust estimate of expected co-occurrence ^d^E_ij_. Following this, we calculated ^d^R_ij_, defined as the ratio of ^d^C_ij_ to ^d^E_ij_ for each species. ^d^R_ij_ is a measure of whether note group j occurs within d-1 notes of note group i more or less often than expected by chance alone. ^d^R_ij_ > 1 suggests note group j occurs within d-1 notes of note group i more often than expected by chance alone, and ^d^R_ij_ < 1 suggests that j occurs within d-1 notes of i less often than expected by chance alone. For the analysis reported in the Results we used a d-value of 4, but also repeated our analysis for d- values of 2 and 6 to ensure that our choice of d did not influence the observed patterns; previous studies also indicate that the choice of d does not change co-occurrence patterns in vocal sequences (Bhat et al. 2022). We chose to report results for a d-value of 4 because, on average, songs of *Spelaeornis* consist of 4-8 notes and a distance of 4 thus captures biologically meaningful note co-occurrence patterns.

To statistically compare transition probability matrices between species, it is necessary that all the Markov chains being compared share the same state space. Therefore, we added a pseudocount of 0.0001 to each entry of the 9×9 transition matrix of each species (thus ensuring that all pairwise transition probabilities were non-zero, and the state space of every species comprised the nine possible note groups defined earlier; for visualizations of transition probabilities we used the original matrices). We then computed the frequency of occurrence for each transition ij in each species’ songs and added this to the pseudocount. This gave us a 9 × 9 transition probability matrix for each species. Next, we employed a homogeneity test that computes a minimum discrimination information statistic (mdis) (Bhat et al. 2022). This statistic tests for differences in the distributions of transition probabilities between samples (in our case species) based on the Kullback-Leibler divergence (Kullback et al. 1962). The test statistic is asymptotically distributed as a ***X*^2^**distribution with ***S*(*S* − 1)(*r* − 1)** degrees of freedom where S is the number of states of the Markov chain (in our case, the number of note groups, i.e., 9) and r is the number of different samples to be compared (in our case, number of species). We computed this statistic for a comparison of all 8 species, as well as pairwise between any two species.

Finally, to test for sequence similarity between species across geographic regions, we computed the pairwise Levenshtein distances between the vocal sequences of each species pair. We then computed a median Levenshtein distance for each species pair and constructed a distance matrix *D*, where *D*_*ab*_ represents the median Levenshtein distance between the song sequences of species *a* and *b*. Using this matrix, we constructed a distance dendrogram using the unweighted pair group method with arithmetic mean (UPGMA). Species closer together on this dendrogram possessed similar vocal sequence structures.

## Results

### Spelaeornis do not exhibit interspecific note parameter divergence

The first three principal components of 10 note parameters explained more than 77% of variation in the notes of *Spelaeornis* (Table S1). PC1 loaded moderately positively on all frequency parameters and loaded negatively on note duration and peak time. PC2 loaded positively on note duration and negatively on minimum peak frequency. PC3 loaded positively on peak time and exhibited weak negative loading on average entropy.

Broadly, *S. ca* and *S. tr* occupy opposite ends of the three-dimensional signal space encompassing all the note parameters of *Spelaeornis*. All other species of *Spelaeornis* heavily overlap with each other in between these extremes (Fig 2B). This suggests that the song notes of *Spelaeornis* have not diverged in allopatry, and that their songs have thus retained similar building blocks across their range. This assertion was also supported by a randomization test (see Methods). We found that the average interspecific distance in note parameter space of *Spelaeornis* is much lower (*Z-*score = -40.248, p < 0.001) than expected by chance, as calculated from 1000 randomized datasets (See Methods). Linear Discriminant Analysis further supports an overlap of notes in signal space. The LDA classifier correctly assigned notes to different species with an accuracy of only 55.8% (Fig S1). The notes of *S. ca* and *S. tr*, occupying opposite edges in *Spelaeornis***’** signal space, were correctly classified at rates of 81.8% and 89.0% respectively. However, these were the only two species with high classification rates: the classification accuracies for all other species were less than 60% (Range: 11.6% for *S. re* – 55.2% for *S. oa*) (Fig S1). Taken together, these statistical and quantitative analyses suggest that allopatric *Spelaeornis* do not exhibit interspecific divergence in song note parameters. Instead, they exhibit more overlap in signal space than expected by chance, consistent with different species using similar notes.

### Note types and note groups in Spelaeornis

We identified multiple note types in each species, and verified these classifications using both linear discriminant classifiers and independent cross-verification by a second author. In total, the number of note types identified were as follows: *S. ca:* 6 note types (1 note type with 2 subclasses, see Methods); *S. ba*: 8 note types; *S. tr:* 21 note types (1 note type with 2 subclasses); *S. ch:* 8 note types; *S. re:* 9 note types; *S. ki:* 6 note types; *S. oa:* 9 note types; *S. lo:* 13 note types (1 note type with 2 subclasses) (Fig S2). Classification accuracies for note types ranged from 71.5% (*S. ca*) to 85.6% (*S. ba*). Additionally, we cross-verified our note type classification through a second observer in a double-blind paradigm. We found an average inter-observer agreement of 76.33% for classification of note types across species.

Because different species of *Spelaeornis* possess similar note parameters, as evinced by heavy overlap in signal space, we grouped notes into note groups for interspecific comparison based on spectral shape (see Methods). We identified the following note groups across species (letters indicate abbreviations for note groups as per the key provided in the Methods):- *S. ca:* a,b,c,d,e,f,g,k; *S. ba:* a,b,c,d,e,f,k; *S. tr:* a,b,c,d,e,f,g,k,l; *S. ch:* a,b,c,d,f; *S. re:* a,b,c,d; *S. ki:* a,b,c,d,f,k; *S. oa:* a,b,c,d,e,f,g; *S. lo:* a,b,c,d,e,f,g,k. The average interobserver agreement in a double-blind note group classification paradigm was 77.75%. It is important to note here that many note groups are shared across species, consistent with our interpretation of the interspecific overlap in signal space. The l note group, however, was unique to *S. tr* and was not observed in other species. Accumulation curves for all species for both note types and note groups were characterized by early saturation, providing evidence that our dataset reliably sampled the note repertoires in *Spelaeornis* (Fig S3, S4).

### Spelaeornis species differ in song complexity

The song complexity of *Spelaeornis* exhibited significant interspecific differences (Kruskal-Wallis test, H=57.05, df=7, p-value= 5.83e-10). Qualitatively, we observed two groups, one consisting of species possessing lower song complexity (fewer note types as a proportion of song length), and the other comprising species with higher song complexity (higher number of note types as a proportion of song length) (Fig 2C). We performed post-hoc pairwise comparisons using Dunn’s Test, and 10 out of 28 species pairs exhibited statistically significant differences in song complexity (Table S2). These differences are represented as compact letter displays (*a-d*) in Fig 2C. *S. ki* differed significantly from *S. oa, S. ch, S. lo* and *S. ba*, all species with higher song complexity (*a*), but not from other species. *S. ca, S. tr* and *S. re* differed significantly from *S. ba* and *S. lo*, but not from *S. oa, S. ch* and *S. ki* (*b*). *S. ch* and *S. oa* differed significantly only from *S. ki (c). S. ba* and *S. lo* significantly differed from *S. ca, S. tr, S. re* and *S. ki (d)*. Thus, in spite of using similar building blocks (i.e. notes) in their songs, species of *Spelaeornis* differ in the complexity of these songs. This suggests that the temporal sequence or syntax of the songs varies between species, a possibility we next investigated using computational analyses of vocal sequence structure.

### Syntactic divergence in the songs of Spelaeornis

We employed a combination of first-order Markov chain analysis, and a note co-occurrence analysis that does not assume Markovianity (see Methods) to examine the syntactic structure of *Spelaeornis* songs. As hinted at by the results of the song complexity analysis, we found that different species fell into three broad syntactic groups. *S. ki* and *S. re*, both occurring in Southeast Asia, exhibited songs with relatively sparse inter-note transitions, with notes instead exhibiting a high tendency to repeat themselves. This was also borne out by the note co-occurrence analysis, where high probabilities of co-occurrence (^d^R_ij_) were primarily along the diagonal of the matrix, indicating that notes tended to co-occur with themselves more than other notes (Fig 3). We defined this sequence structure as Repetitive syntax. On the other hand, the songs of *S. ca* from the Eastern Himalayas consisted of 2 or 3 notes alternating with each other. This is represented by the reciprocating arrows between different nodes in Fig 3, as well as the note co-occurrence matrix. Here, higher probabilities occurred both along the diagonal and with 1-2 other notes, indicating that each note type co-occurred with itself and one or two other notes. We defined this second syntactic type as Alternating syntax. Finally, we also observed songs with Complex syntax, primarily exhibited by *S. oa, S. lo*, and *S. ch* which occur in the hills south of the Brahmaputra River. In these species’ songs, each note group exhibited transitions to many different note groups, manifested as many arrows originating from each node in Fig 4. In the co-occurrence matrices, probabilities were generally low, and did not show any pattern of higher values either on- or off-diagonal, indicating that each note could generally co-occur with many other note types.

**Fig 3:**
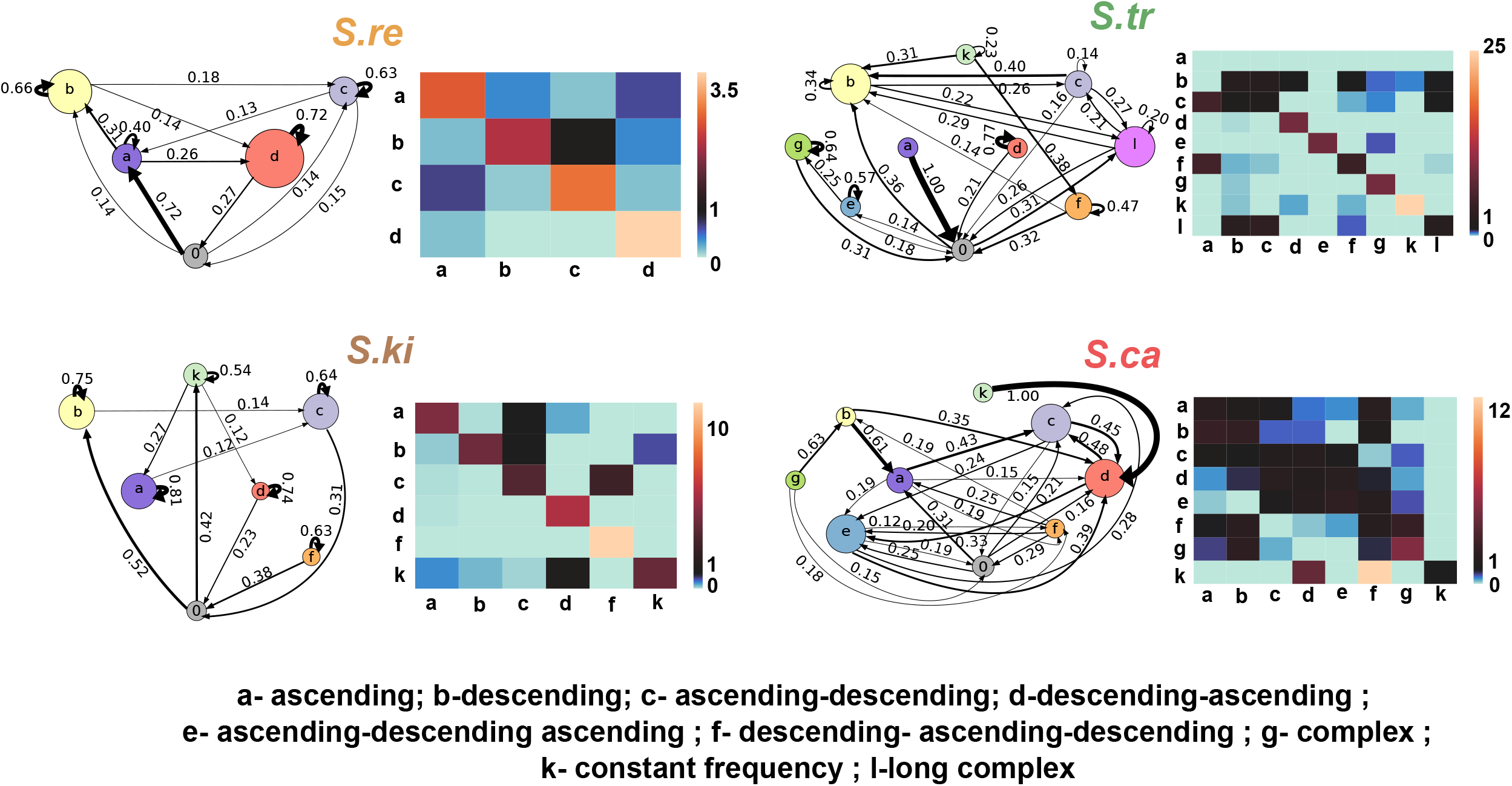
Transition probabilities obtained by modeling note group sequences of *Spelaeornis* as first-order Markov chains (left side on all the panels). Different colors represent different note groups (see key below the figure), and the size of the node is proportional to frequency of occurrence in our dataset. The thickness of each arrow scales with increasing transition probabilities between note groups. Ratio of observed to expected probability of co-occurrences for note groups for a value of d=3 (right side on all the panels). Warmer or redder colors in the ***ij***-th square on the matrix depict that note group i occurs with note group j more often than by chance. *S. ki* and *S. re* exhibit repetitive syntax, depicted by thicker arrows for self-transitions and redder colors along the diagonal in the co-occurrence matrix. In *S. ca*, we find alternating syntax, depicted by arrows going back and forth between different note groups and redder colors both above and below the diagonal in the co-occurrence matrix. *S. tr* exhibits a combination of alternating and repetitive syntax. Some note groups exhibit higher propensity for self-transition in the transition probability figures and redder colors along the diagonal in the co- occurrence matrix, whereas others exhibit alternation (redder colors for 1-2 other note groups in the co-occurrence matrix).

**Fig 4:**
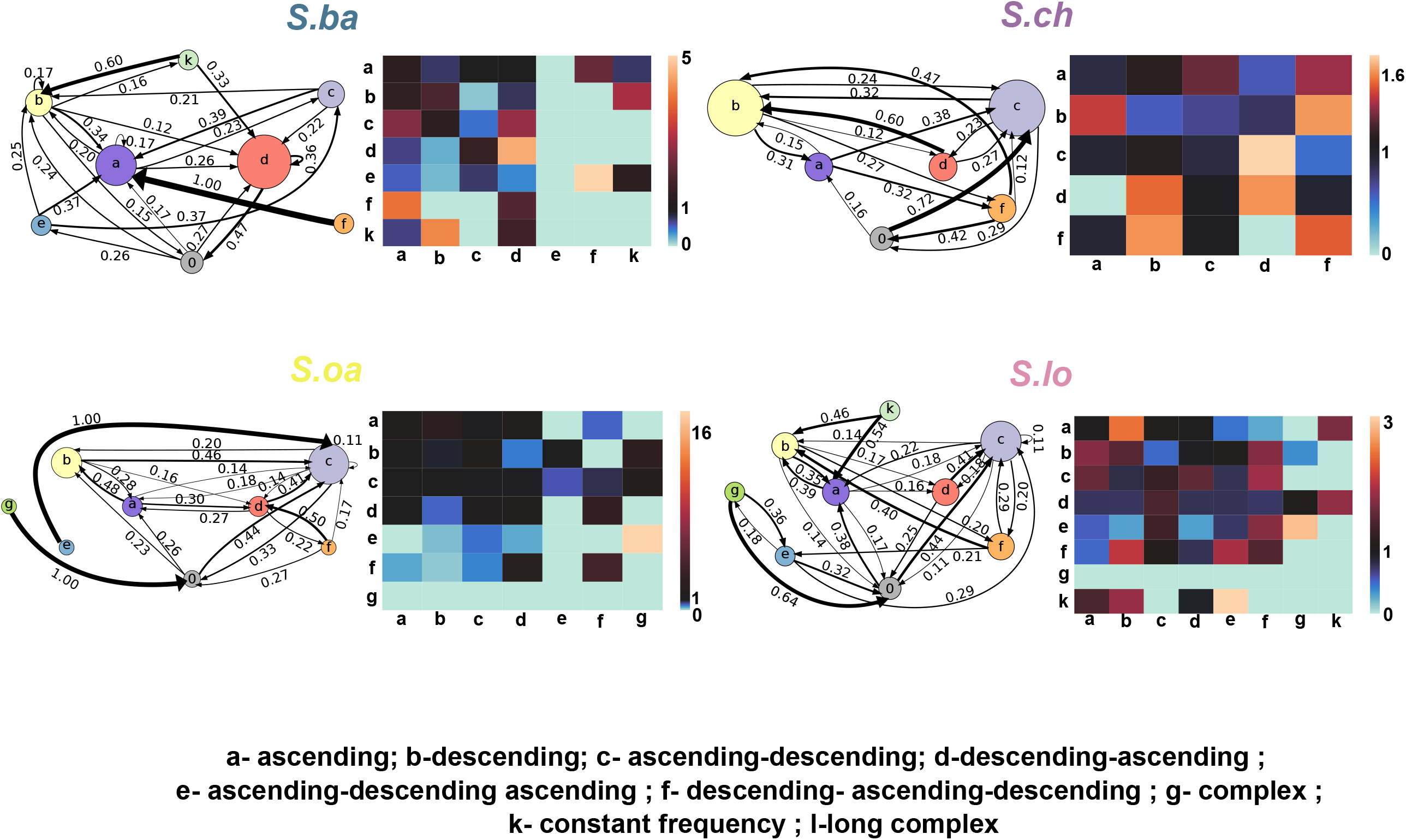
Transition probabilities and the ratio of observed to expected probability of co- occurrences for note groups for a value of d=3, as in Fig 3. *S. lo, S. oa* and *S. ch* exhibit complex syntax, where each group can transition to many others with a low probability of self- transition and a lack of discernible structure in the co-occurrence matrix. *S. ba*, on the other hand, exhibits an intermediate between complex and repetitive syntax. Many of the above features are present, but some note groups exhibit an increased propensity to self-transition, resulting in some repetition within vocal sequences (see Supplementary).

Two of the eight species (*S. ba* and *S. tr*) in the genus exhibited evidence of intermediate syntactic types, employing a combination of the syntactic rules outlined above to order notes. *S. ba* songs suggested an intermediate between Repetitive and Complex Syntax. As depicted in Fig 4, some notes have high probabilities of transition to many different note groups in addition to having a relatively high probability of self-transitions (i.e., repetitions). This is also observed in the co-occurrence matrix, where generally low values of ^d^R_ij_ are coupled with higher values along the diagonal for certain notes. An examination of the song sequences (Supplementary File 1) suggested that these two types could occur within a single song, with multiple note groups in a single song, some of them repeating (unlike the other species with complex syntax). On the other hand, *S. tr* appears to use a combination of Alternating and Repetitive syntax. Both the transition probability matrix and the note co-occurrence matrix confirm this, with some note groups tending to repeat and others alternating between each other. The song sequences indicate that unlike *S. ba, S. tr* uses either alternating or repetitive syntax in a song, and not a combination of both (Supplementary File 2). This indicates the presence of different song types with differing syntax within this species, although we did not have enough data to determine if these followed a geographic pattern. The patterns described above are broadly consistent across different values of d (Fig S5, S6), in keeping with the findings of other studies that have used the same analysis (Bhat et al. 2022). Similar patterns were also observed when considering note types instead of note groups (Fig S7), which, coupled with our finding of overlap in note parameter space, suggests that species of *Spelaeornis* use similar notes, but arrange them according to different syntactic rules. Our note group classification thus represents a valid means of interspecific comparison.

Using the aforementioned homogeneity test for Markov chains, we found that the transition probability matrices for different species of *Spelaeornis* differ significantly from each other (***X*^2^** = 8061.23, df=504, p-value=0). Pairwise species comparisons found that transition probability matrices differ for all species pairs (Fig S8). Further, we also found that species exhibiting similar syntactic structure generally tend to have a smaller value of the test statistic when compared to species with different syntactic rules (Fig S8). For example, *S. ki* and *S. re* were closest to each other (smaller values of the test statistic), whereas *S. ch, S. oa* and *S. lo* were closer to each other than to other species. *S. ca* was relatively far apart from all other species (higher values of the test statistic). *S. ba* and *S. tr* were closest to *S. re*, and *S. tr* was also close to *S. ki* (all four species use at least some repetitive syntax), but neither *S. ba* nor *S. tr* were particularly close to any of the other species.

### Syntactic structure of vocal sequences follows geographic barriers

Our analyses suggested that species of *Spelaeornis* existing geographically closer to each other possess similar syntactic structure in their songs. Although the proportions of different note groups in the sequences analyzed differed across geographically proximate species (Fig 5A), the syntactic rules employed to construct songs were conserved within each sub-region. However, across sub-regions separated by geographic barriers, syntactic structures changed altogether (Fig 3, 4). We thus predicted that the vocal sequences of geographically proximate species should group together, i.e. exhibit greater similarity than those from different geographic regions. To further quantitatively examine sequence similarity across species, we calculated sequence similarity using the median interspecific Levenshtein Distance and constructed a distance dendrogram, using the UPGMA method (see Methods). We observed that species geographically closer to each other are also closer to each other on the distance dendrogram (Fig 5B), supporting the idea that geographic barriers drive syntactic diversification in the absence of note divergence. To illustrate, *S. ki* and *S. re*, found in South-East Asia and employing repetitive syntax grouped together in the dendrogram (Fig 5B). Additionally, *S. oa, S. ch*, and *S. lo*, found south of the Brahmaputra River in North-Eastern India and employing complex syntax, formed a second group in the dendrogram (Fig 5B), close to *S. ba* which occupies the junction of the hills north and south of the region (and exhibits an intermediate syntax with elements of complex syntax, see above). Taken together, our results suggest that in the absence of note divergence, acoustic signal diversification in *Spelaeornis* involves changes in the syntactic rules employed to order sequences across geographic barriers. Thus, vocal sequences of species that are geographically proximate tend to structurally resemble each other, whereas species from different geographic regions exhibit distinct vocal sequence structures.

**Fig 5:**
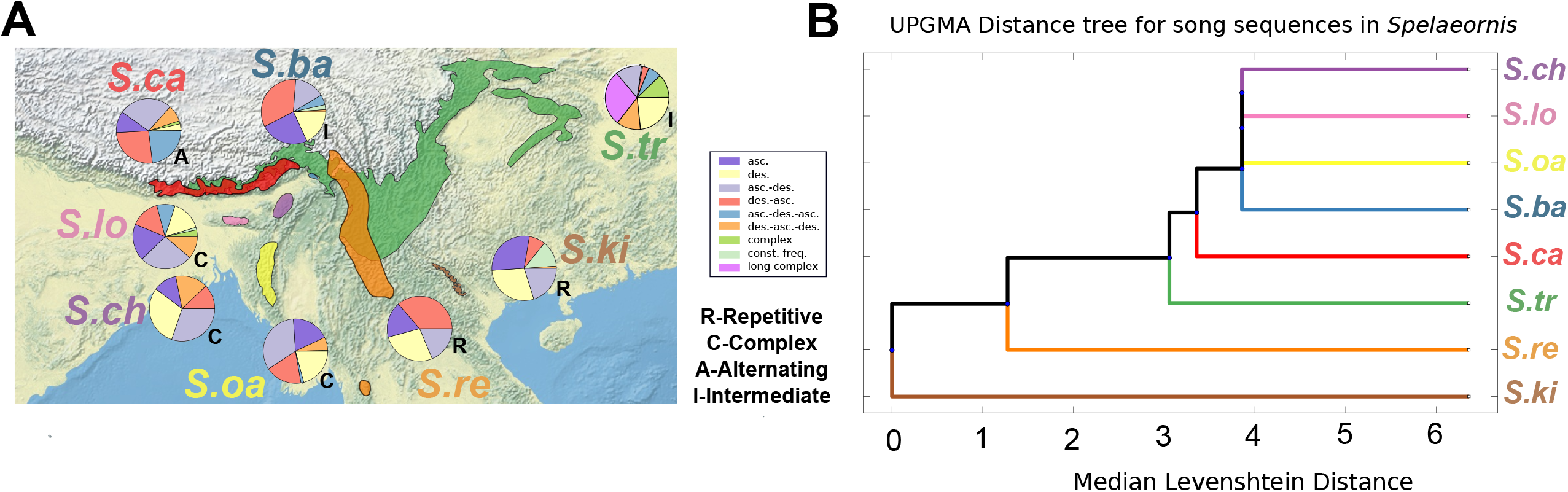
**A**. Pie charts representing the proportions of different note groups across species in our dataset. Note accumulation curves suggest that this dataset is an adequate sample of the note group repertoire across species (see Fig S3). Species geographically closer to each other exhibit similar syntactic structure (see alphabets), with changes across major geographic barriers. Note in particular the intermediate syntax of *S. ba*, which occupies a junction between two major geographic regions. **B**. UPGMA distance dendrogram constructed using median Levenshtein distances between vocal sequences for each pair of species. Species with similar vocal sequences cluster together in this distance dendrogram, and are also geographically proximate, as shown in the map.

## Discussion

In summary, we uncover evidence that song notes in the genus *Spelaeornis* remain more or less conserved across species (Fig 2B, S1), but are arranged into songs according to different syntactic rules (Fig 3, 4) based on the presence of geographic barriers. We found three broad syntactic groups (with two species exhibiting intermediate syntax), occupying the Eastern Himalayas, the hills south of the Brahmaputra river, and Southeast Asia respectively (Fig 3, 4). Within each of these regions, geographically proximate species exhibited similar syntactic structure in their songs (supported by the Kullback homogeneity test, and by the UPGMA distance dendrogram) (Fig 5B, S8), with only minor differences in the proportions of different note groups within their vocal sequences (Fig 5A). Our analysis of note accumulation suggested that our sampling of the note group repertoires for each species is adequate (Fig S3), and therefore we hypothesize that our results represent genuine interspecific changes in syntactic structure of vocal sequences across geographic barriers. Because a variety of methods suggest that we have adequately sampled the note group repertoire in this genus, we believe that additional recordings will not significantly alter the pattern we observed, particularly that of groups across biogeographic regions defined by vocal sequence structure.

Previous studies on interspecific song divergence have largely focused on differences in note parameters, or on broad, summarized metrics of song structure such as trill rate or song duration (Slabbekoorn and Smith 2002a; Slabbekoorn and Smith 2002b; Haavie et al. 2004; Podos and Warren 2007; Irwin et al. 2008; Grant and Grant 2010; Tobias et al. 2010; Podos et al. 2013; Wilkins et al. 2013). Here, employing a combination of tools (Kershenbaum et al. 2012), including a robust computational analysis of note co-occurrence originally developed to study syntactic structure in anuran vocalizations (Bhat et al. 2022), we demonstrate that *Spelaeornis* represents an example of syntactic diversification. This implies that the rules by which notes are temporally arranged into songs have diversified across geographic barriers without significant change in the underlying notes themselves. Our statistically robust analytical framework enables us to comparatively examine temporal structure in diverse animal signals and thus investigate signal evolution at multiple levels, from individual notes to syntax.

Contrary to our initial predictions, we did not find any interspecific divergence in the note parameters in *Spelaeornis*. Instead, the overlap in signal space as well as a statistical randomization test suggest that *Spelaeornis* species use similar song notes. In light of our findings that species geographically closer to each other exhibit similar syntax (Fig 5B), and also share note groups (Fig 2B), future research should comparatively examine the role of notes versus syntax in species recognition. This will be particularly interesting for the case of *S. badeigularis*, which occupies a junction point between different biogeographic regions, and also exhibits songs with elements of both repetitive and complex syntax. Finally, the high-altitude *S. troglodytoides* exhibits distinct song types with alternating and repetitive syntax, and more data is required to understand whether this variation also corresponds to geographic barriers at high altitudes. Distinct song types with different syntactic structures may also occur as rarer songs in other species, particularly those such as *S. badeigularis* that occupy junctions between geographic regions (King and Donahue 2006). However, the addition of rare song types is unlikely to significantly alter the salient pattern we observe, that of syntactic change across geographic regions without change in the underlying song notes. Therefore, the syntactic types we describe here likely represent the dominant forms within each geographic region. *Spelaeornis* as a genus is very poorly studied, with some species going undetected for decades before being rediscovered (King and Donahue 2006). In light of this, online song databases provide us with valuable data which, coupled with a rigorous analytical framework, enables us to effectively examine questions about behavior and signal evolution. Quantitative analysis (Fig S3, S4) suggests that we have adequately sampled the note group and note type repertoire of all species within the genus (including for *S. reptatus* for which we had the smallest sample size), further supporting this assertion. With so little known about the different hill ranges in the Himalayas and Southeast Asia, it is possible that undescribed species in this genus remain to be found, and their songs may prove valuable in understanding the steps by which syntactic diversification of bird song has occurred.

The evolution and function of bird song exhibits many parallels to human languages (Doupe and Kuhl 1999; Salwiczek and Wickler 2004; Berwick et al. 2011; Miyagawa et al. 2013; Collier et al. 2014; Sainburg et al. 2019), and geography shapes the structure and syntax of human languages as well (Henry et al. 2015; Dunn 2019; Huisman et al. 2019; Urban 2021). It is, therefore, also interesting to note that diversification of human languages in the region follows a similar geographic pattern (Sagart et al. 2019), with an Indo-Chinese trail of language diversification north of the Brahmaputra River and the Burman language groups diversifying south of the Brahmaputra River. The parallels in the evolution of diverse culturally transmitted traits point to the importance of studying syntax and sequences in birdsong to gain broader evolutionary insights.

Acoustic signals in general are shaped by various factors such as evolutionary history, geographic separation, and sexual selection (Marler and Tamura 1962; McCracken and Sheldon 1997; Baker 2006; Baker et al. 2006; Podos and Warren 2007; Kirschel et al. 2009; Kershenbaum et al. 2012; Garland et al. 2013; Lachlan et al. 2013; Seddon et al. 2013; Wilkins et al. 2013; Lachlan et al. 2016; Arato and Fitch 2021). By presenting evidence that a) despite geographical isolation, *Spelaeornis* appear to possess similar song notes across species, and b) syntactic structure of vocal sequences in *Spelaeornis* has diverged across geographical barriers, we shed light on the diverse paths through which signals evolve in allopatry (Kershenbaum et al. 2012). Because, as discussed above, bird song exhibits similar geographic patterns to those observed for human languages, we suggest that the computational approach employed here can have far-reaching implications in our understanding of speciation, signal evolution and linguistic diversification. The Himalayas and South-east Asia are regions with hyperdiverse ecological communities (Srinivasan et al. 2014; Mungee and Athreya 2021; Ashokan et al. 2022). Studies such as ours emphasize the importance of studying biodiversity, natural history and behavior of taxa in these regions to understand the biogeography of communication strategies and signal evolution.

## Acknowledgments and Funding

We are grateful to the many recordists (listed in the Supplementary), whose recordings from public databases contributed to this project, as well as to Ramana Athreya, Umesh Srinivasan, Vaibhav Chhaya, Aurnab Ghose and members of the Krishnan lab for useful feedback and discussions. Finally, we thank Seshadri KS for help with maps. AK is funded by an INSPIRE Faculty Award from the Department of Science and Technology, Government of India, and an Early Career Research Grant (ECR/2017/001527) from the Science and Engineering Research Board (SERB), Government of India. MAJ and ASB are the recipients of the KVPY Fellowship from the Government of India.

---

**Fig S1:** Interspecific confusion matrix representing correct classification rates of the note parameters of different species in a linear discriminant classifier built using MATLAB, with the 10 note parameters described above as the input. Percentages in the blue boxes indicate correct classification rates for each species, and the red boxes indicate misclassification rates. We observe an overall classification accuracy of 55.8%, coupled with high misclassification rates of LDA in assigning notes to species.

**Fig S2:** Intraspecific confusion matrices representing correct classification rates for individual note types of each species using linear discriminant classifiers built in MATLAB.

**Fig S3:** The accumulation curves for note groups exhibit early saturation, suggesting that we adequately sampled the note groups for each species.

**Fig S4:** The accumulation curves for note types exhibit early saturation, suggesting that we adequately sampled the note types for each species.

**Fig S5:** Ratio of observed to expected probability of co-occurrences for note groups for a value of d=2.

**Fig S6:** Ratio of observed to expected probability of co-occurrences for note groups for a value of d=6.

**Fig S7:** Transition probabilities obtained by modeling note type sequences of *Spelaeornis* as first-order Markov chains (left side on all the panels), as in Fig 3 and 4 (which depict the same analysis for note groups). Ratio of observed to expected probability of co-occurrences for note types for a value of d=3 (right side on all the panels), as in Fig 3 and 4. The syntactic patterns described for note groups in *Spelaeornis* remain consistent when considering note types. Thus, interspecific syntactic patterns observed are not influenced by our note group classification.

**Fig S8:** Heat map depicting the pairwise test statistic of the homogeneity test. Species geographically closer to each other generally exhibit a smaller value of the test statistic, suggesting they are more similar to each other than to other congeners.

**Table S1:** Results of Principal Component analysis on the correlation matrix of the ten note parameters measured from the songs of *Spelaeornis*. The values represent the loadings of various parameters onto the Principal Components (PCs), as well as the proportion of explained variance.

**Table S2:** Table depicting the results of post-hoc Dunn’s tests for pairwise comparisons. Species pairs showing statistically significant differences in song complexity are represented in bold.

