## Supplementary Figures for "Allopatric montane wren-babblers exhibit similar song notes but divergent vocal sequences"

Fig S1

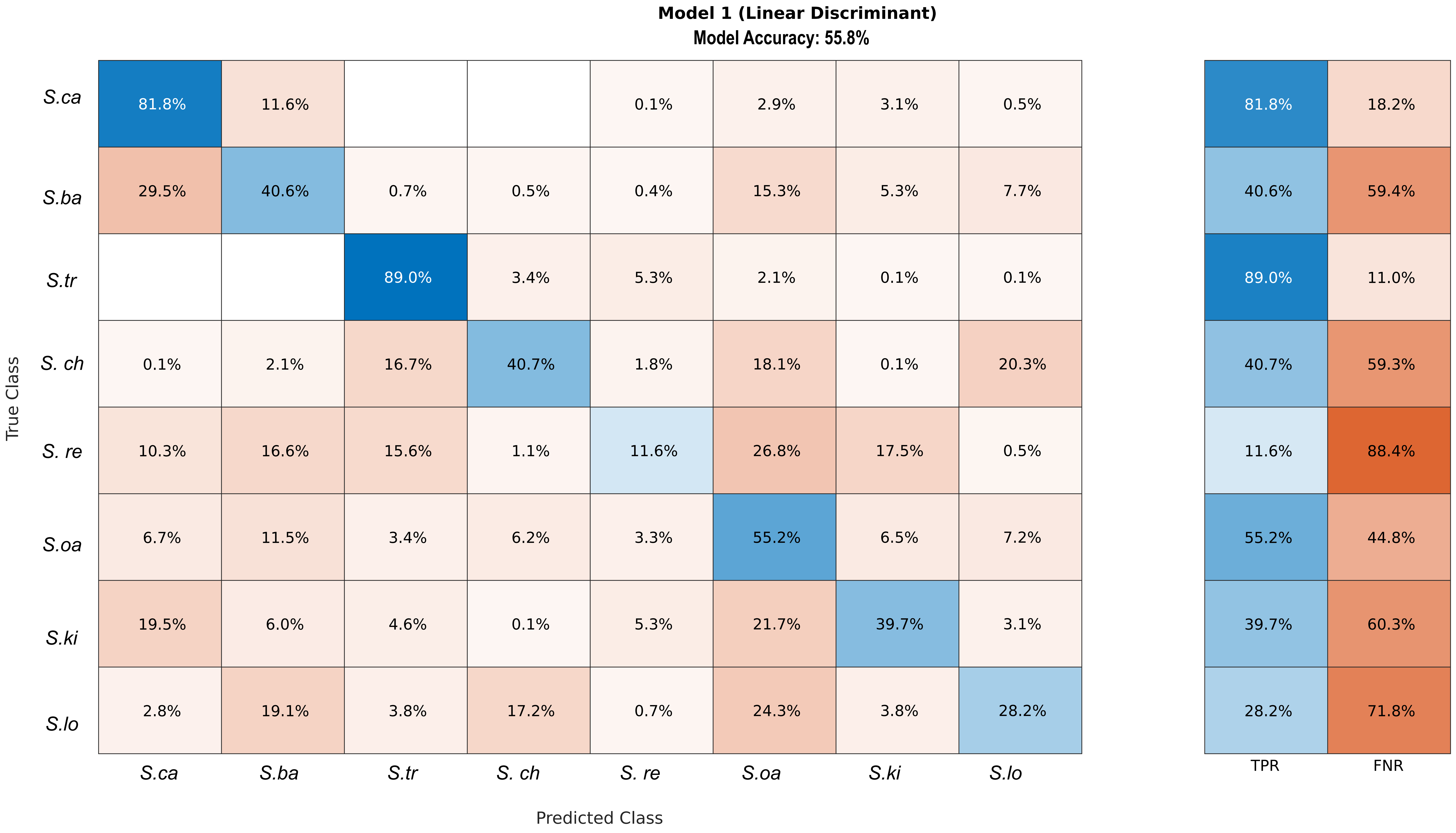
Model 1 (Linear Discriminant)
Model Accuracy: 55.8%
True Class
Predicted Class
S.ca S.ba S.tr S. ch S. re S.oa S.ki S.lo
TPR FNR
S.ca: 81.8% 11.6% 0.1% 2.9% 3.1% 0.5% | 81.8% 18.2%
S.ba: 29.5% 40.6% 0.7% 0.5% 0.4% 15.3% 5.3% 7.7% | 40.6% 59.4%
S.tr: 89.0% 3.4% 5.3% 2.1% 0.1% 0.1% | 89.0% 11.0%
S. ch: 0.1% 2.1% 16.7% 40.7% 1.8% 18.1% 0.1% 20.3% | 40.7% 59.3%
S. re: 10.3% 16.6% 15.6% 1.1% 11.6% 26.8% 17.5% 0.5% | 11.6% 88.4%
S.oa: 6.7% 11.5% 3.4% 6.2% 3.3% 55.2% 6.5% 7.2% | 55.2% 44.8%
S.ki: 19.5% 6.0% 4.6% 0.1% 5.3% 21.7% 39.7% 3.1% | 39.7% 60.3%
S.lo: 2.8% 19.1% 3.8% 17.2% 0.7% 24.3% 3.8% 28.2% | 28.2% 71.8%

Fig S2

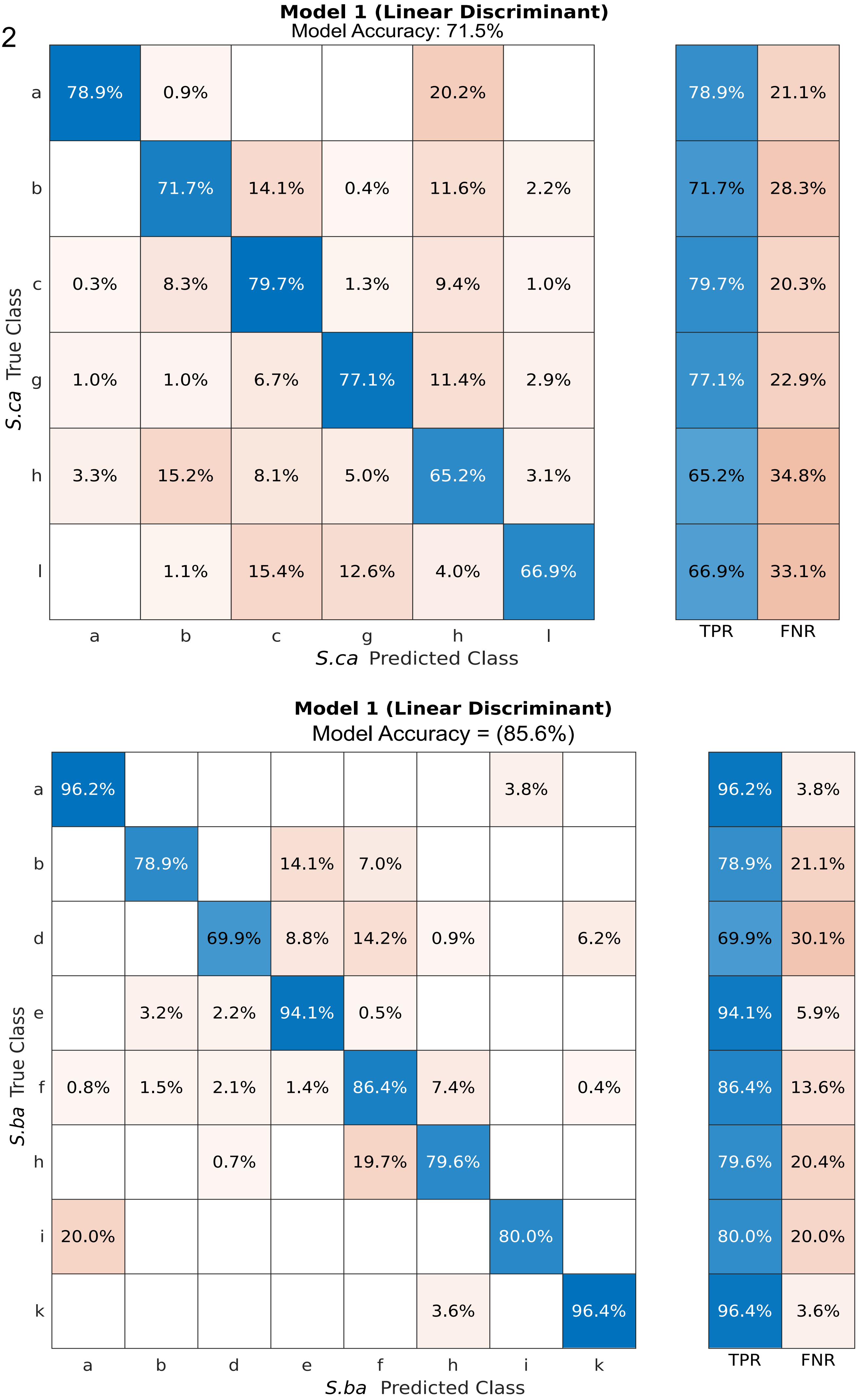
Model 1 (Linear Discriminant)
Model Accuracy: 71.5%
a 78.9% 0.9% 20.2%
b 71.7% 14.1% 0.4% 11.6% 2.2%
c 0.3% 8.3% 79.7% 1.3% 9.4% 1.0%
g 1.0% 1.0% 6.7% 77.1% 11.4% 2.9%
h 3.3% 15.2% 8.1% 5.0% 65.2% 3.1%
l 1.1% 15.4% 12.6% 4.0% 66.9%
a b c g h l
S.ca Predicted Class
S.ca True Class
78.9% 21.1%
71.7% 28.3%
79.7% 20.3%
77.1% 22.9%
65.2% 34.8%
66.9% 33.1%
TPR FNR
Model 1 (Linear Discriminant)
Model Accuracy = (85.6%)
a 96.2% 3.8%
b 78.9% 14.1% 7.0%
d 69.9% 8.8% 14.2% 0.9% 6.2%
e 3.2% 2.2% 94.1% 0.5%
f 0.8% 1.5% 2.1% 1.4% 86.4% 7.4% 0.4%
h 0.7% 19.7% 79.6%
i 20.0% 80.0%
k 3.6% 96.4%
a b d e f h i k
S.ba Predicted Class
S.ba True Class
96.2% 3.8%
78.9% 21.1%
69.9% 30.1%
94.1% 5.9%
86.4% 13.6%
79.6% 20.4%
80.0% 20.0%
96.4% 3.6%
TPR FNR

Fig S2

**Model 1 (Linear Discriminant)**

Model Accuracy 71.8%

| *S.oa* True Class \ *S.oa* Predicted Class | a | b | c | d | g | h | j | k | q | TPR | FNR |
|---|---|---|---|---|---|---|---|---|---|---|---|
| a | 74.1% | 0.4% | 4.4% | | | 7.5% | 2.6% | 0.9% | 10.1% | 74.1% | 25.9% |
| b | | 87.7% | | 3.8% | | | | 7.5% | 0.9% | 87.7% | 12.3% |
| c | 12.7% | 1.1% | 61.1% | 1.4% | 0.8% | 0.6% | 17.5% | 3.7% | 1.1% | 61.1% | 38.9% |
| d | | 4.3% | 26.1% | 69.6% | | | | | | 69.6% | 30.4% |
| g | | | 23.1% | | 76.9% | | | | | 76.9% | 23.1% |
| h | 15.5% | | | | | 84.5% | | | | 84.5% | 15.5% |
| j | 7.1% | | 1.6% | | | 5.8% | 85.5% | | | 85.5% | 14.5% |
| k | 0.5% | 9.8% | 0.5% | 1.0% | | 1.0% | | 59.3% | 27.8% | 59.3% | 40.7% |
| q | 19.2% | 2.9% | | | 4.1% | | 0.6% | 14.5% | 58.7% | 58.7% | 41.3% |

**Model 1 (Linear Discriminant)**

Model Accuracy 79.3%

| *S.lo* True Class \ *S.lo* Predicted Class | a | b | c | d | g | i | k | m | n | o | p | t | u | TPR | FNR |
|---|---|---|---|---|---|---|---|---|---|---|---|---|---|---|---|
| a | 86.7% | | | | 3.3% | 10.0% | | | | | | | | 86.7% | 13.3% |
| b | | 81.7% | 9.9% | | 1.1% | 1.5% | | | 4.9% | | 0.8% | | | 81.7% | 18.3% |
| c | | 18.9% | 70.5% | | 0.8% | | | 1.6% | 8.2% | | | | | 70.5% | 29.5% |
| d | 1.8% | 0.9% | | 77.9% | 5.3% | 3.5% | | 0.9% | 7.1% | | | | 2.7% | 77.9% | 22.1% |
| g | | 3.8% | | | 77.4% | 18.9% | | | | | | | | 77.4% | 22.6% |
| i | 3.4% | 3.4% | | 0.7% | 9.5% | 82.4% | | | 0.7% | | | | | 82.4% | 17.6% |
| k | | 9.1% | | | | | 90.9% | | | | | | | 90.9% | 9.1% |
| m | | 5.0% | | | | | | 75.0% | 20.0% | | | | | 75.0% | 25.0% |
| n | 1.3% | 3.2% | | 2.6% | 2.6% | 7.1% | | 4.5% | 76.6% | | 1.9% | | | 76.6% | 23.4% |
| o | | | | 12.5% | | | | | | 87.5% | | | | 87.5% | 12.5% |
| p | | | | | | | | | | | 100.0% | | | 100.0% | |
| t | | 12.5% | | | | | | | | | | 87.5% | | 87.5% | 12.5% |
| u | | | | | | | | | | | | | 100.0% | 100.0% | |

Fig S2

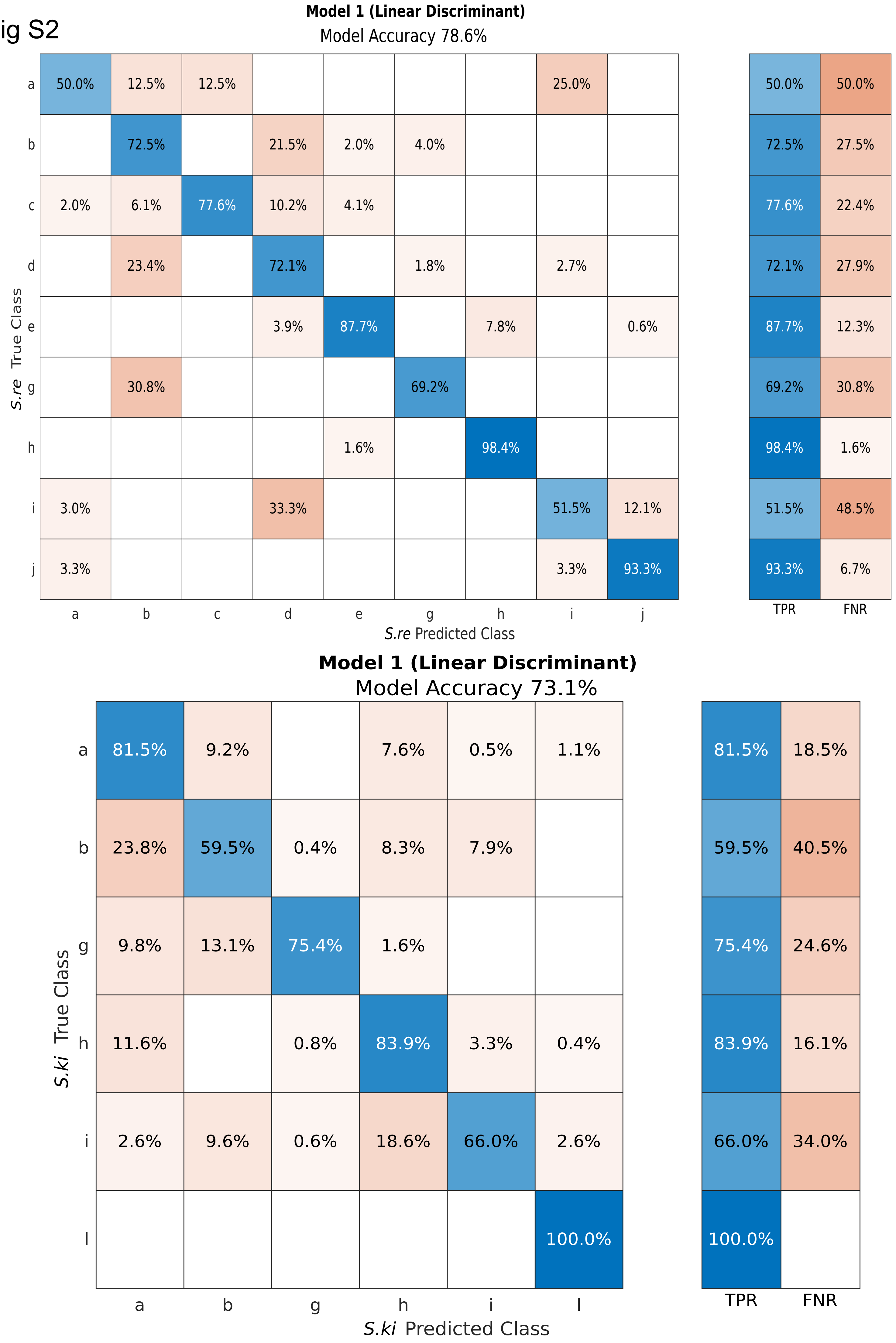

Model 1 (Linear Discriminant)
Model Accuracy 78.6%
S.re True Class
S.re Predicted Class
a b c d e g h i j
TPR FNR
a: 50.0% 12.5% 12.5% 25.0% | 50.0% 50.0%
b: 72.5% 21.5% 2.0% 4.0% | 72.5% 27.5%
c: 2.0% 6.1% 77.6% 10.2% 4.1% | 77.6% 22.4%
d: 23.4% 72.1% 1.8% 2.7% | 72.1% 27.9%
e: 3.9% 87.7% 7.8% 0.6% | 87.7% 12.3%
g: 30.8% 69.2% | 69.2% 30.8%
h: 1.6% 98.4% | 98.4% 1.6%
i: 3.0% 33.3% 51.5% 12.1% | 51.5% 48.5%
j: 3.3% 3.3% 93.3% | 93.3% 6.7%
Model 1 (Linear Discriminant)
Model Accuracy 73.1%
S.ki True Class
S.ki Predicted Class
a b g h i l
TPR FNR
a: 81.5% 9.2% 7.6% 0.5% 1.1% | 81.5% 18.5%
b: 23.8% 59.5% 0.4% 8.3% 7.9% | 59.5% 40.5%
g: 9.8% 13.1% 75.4% 1.6% | 75.4% 24.6%
h: 11.6% 0.8% 83.9% 3.3% 0.4% | 83.9% 16.1%
i: 2.6% 9.6% 0.6% 18.6% 66.0% 2.6% | 66.0% 34.0%
l: 100.0% | 100.0%

Fig S2

**Model 1 (Linear Discriminant)**

Model Accuracy: 77.3%

| *S.ch* True Class \ *S.ch* Predicted Class | b | c | d | i | l | q | r | u | TPR | FNR |
|---|---|---|---|---|---|---|---|---|---|---|
| b | 95.9% | | | | | 4.1% | | | 95.9% | 4.1% |
| c | 4.0% | 65.2% | 9.1% | 0.4% | 2.2% | 12.3% | 0.7% | 6.2% | 65.2% | 34.8% |
| d | 4.8% | 17.7% | 67.7% | | 6.5% | 3.2% | | | 67.7% | 32.3% |
| i | | 31.2% | | 68.8% | | | | | 68.8% | 31.2% |
| l | | 5.3% | 7.9% | 2.6% | 84.2% | | | | 84.2% | 15.8% |
| q | | 12.1% | 0.4% | 0.4% | | 86.2% | 0.8% | | 86.2% | 13.8% |
| r | | | | | | 25.0% | 75.0% | | 75.0% | 25.0% |
| u | | 5.6% | | | | | | 94.4% | 94.4% | 5.6% |

**Model 1 (Linear Discriminant)**

Model Accuracy 84.7%

| *S.tr* True Class \ *S.tr* Predicted Class | b | b1 | c1 | d | d1 | e | f | f1 | h | h1 | i1 | k | m | p | q | s | u | w | x | y | z | | |
|---|---|---|---|---|---|---|---|---|---|---|---|---|---|---|---|---|---|---|---|---|---|---|---|
| b | 73.9% | | | 14.5% | | | | | | | 1.4% | | 2.9% | | | | 1.4% | 1.4% | 2.9% | 1.4% | | 73.9% | 26.1% |
| b1 | | 100.0% | | | | | | | | | | | | | | | | | | | | 100.0% | |
| c1 | | | 94.1% | | | | | | | | | | | 2.5% | | | 1.7% | 0.8% | | 0.8% | | 94.1% | 5.9% |
| d | 6.7% | | | 80.0% | | | | | | | | | | | | | | 13.3% | | | | 80.0% | 20.0% |
| d1 | | | | | 100.0% | | | | | | | | | | | | | | | | | 100.0% | |
| e | | | | | | 74.2% | | 6.5% | | | 3.2% | 9.7% | | | | | | | 3.2% | 3.2% | | 74.2% | 25.8% |
| f | | | | | | | 100.0% | | | | | | | | | | | | | | | 100.0% | |
| f1 | | | | | | | | 89.1% | | | | 10.9% | | | | | | | | | | 89.1% | 10.9% |
| h | | | 0.6% | | | | | | 69.2% | | | 29.6% | | | | | | 0.6% | | | | 69.2% | 30.8% |
| h1 | | | | | | | | | | 100.0% | | | | | | | | | | | | 100.0% | |
| i1 | | | | | | | | | | | 80.0% | | | | | 20.0% | | | | | | 80.0% | 20.0% |
| k | 0.5% | | 0.3% | 0.3% | | 0.2% | 0.2% | 0.2% | 4.6% | | | 90.3% | | | | | 0.7% | 0.7% | | 1.2% | 1.0% | 90.3% | 9.7% |
| m | | | | | | | | | | | | | 100.0% | | | | | | | | | 100.0% | |
| p | | | 5.6% | | | | | | | | | | | 94.4% | | | | | | | | 94.4% | 5.6% |
| q | 3.7% | | | | | | | | | | | | | | 96.3% | | | | | | | 96.3% | 3.7% |
| s | | 0.8% | | | | | | | | | | | | | | 90.1% | | | | 9.1% | | 90.1% | 9.9% |
| u | | 8.3% | 6.7% | | | | | 5.0% | | | | | 3.3% | 3.3% | | | 63.3% | 1.7% | | 8.3% | | 63.3% | 36.7% |
| w | | | | | | | | | | | | 5.3% | | | | | | 94.7% | | | | 94.7% | 5.3% |
| x | | | | | | | | 1.7% | | | | | 3.4% | | | | | | 94.8% | | | 94.8% | 5.2% |
| y | 0.8% | | | | | | | 4.2% | | | | 0.8% | | | | | 6.7% | | | 87.4% | | 87.4% | 12.6% |
| z | 2.5% | | | | 7.5% | | | | | | | | | | | | 12.5% | | | 5.0% | 72.5% | 72.5% | 27.5% |

Fig S3

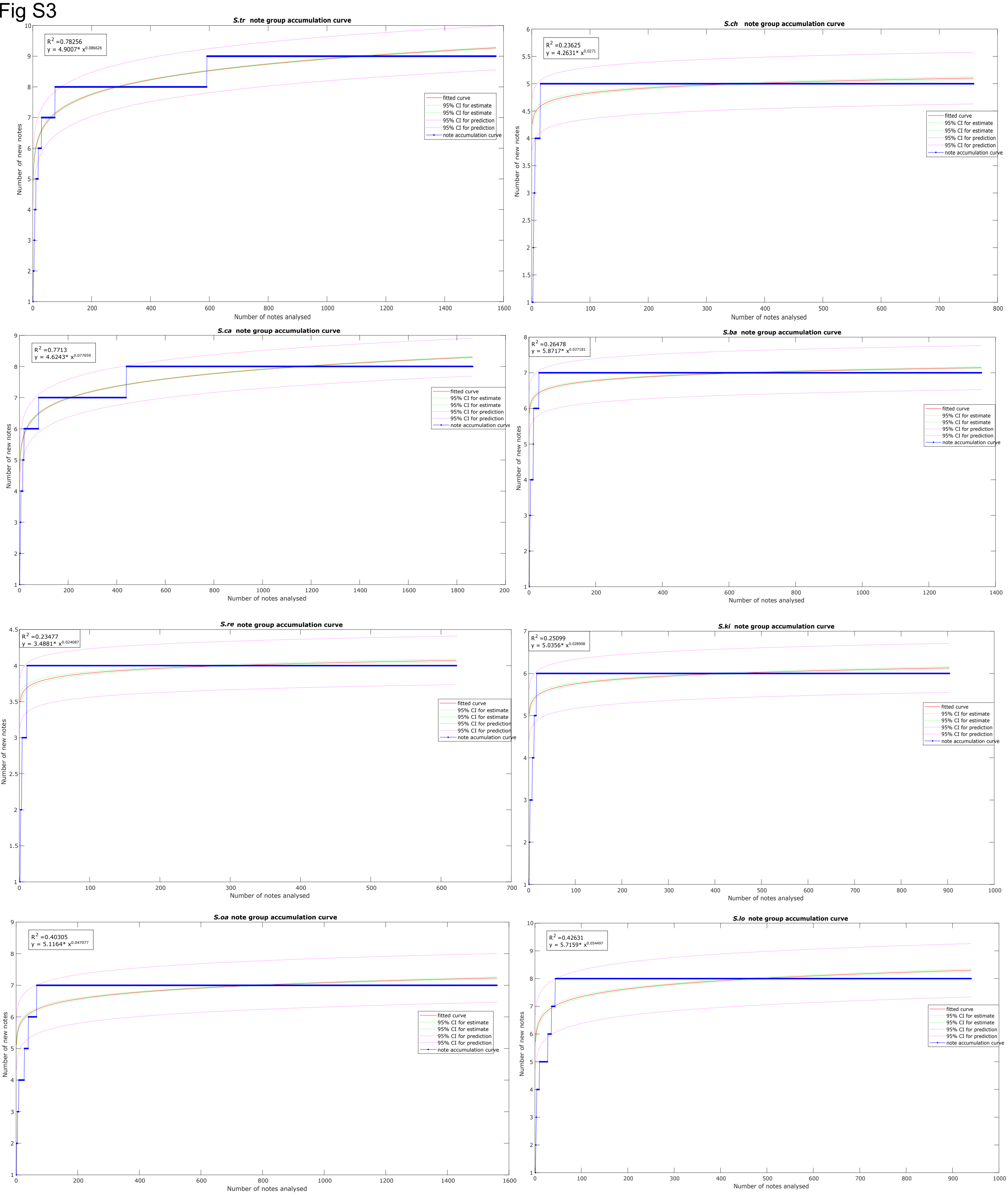


Fig S3: The accumulation curves for note groups exhibit early saturation, suggesting that we adequately sampled all the note groups for each species.

Fig S4

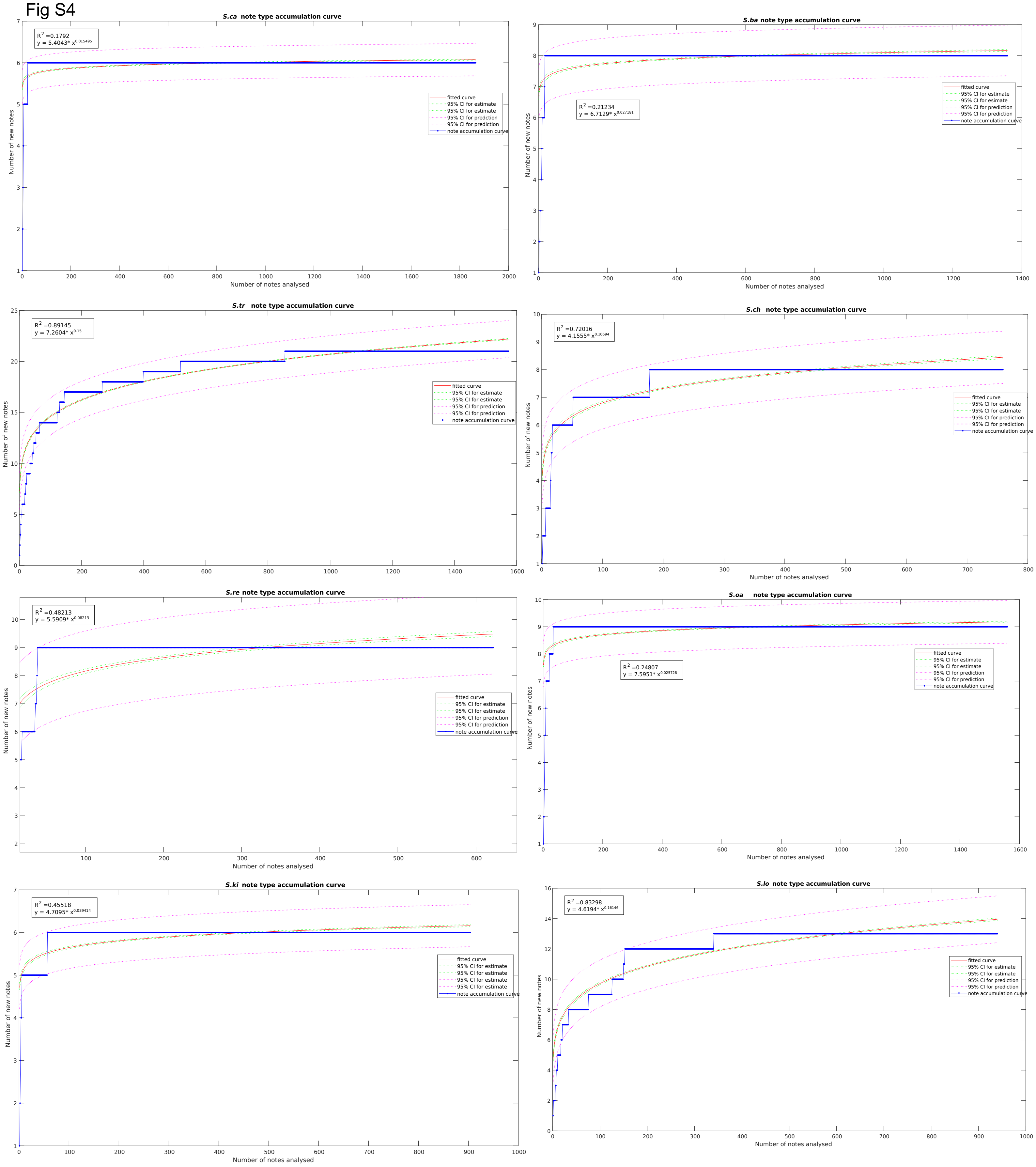


Fig S4: The accumulation curves for note types exhibit early saturation, suggesting that we adequately sampled all the note types for each species.

Fig S5

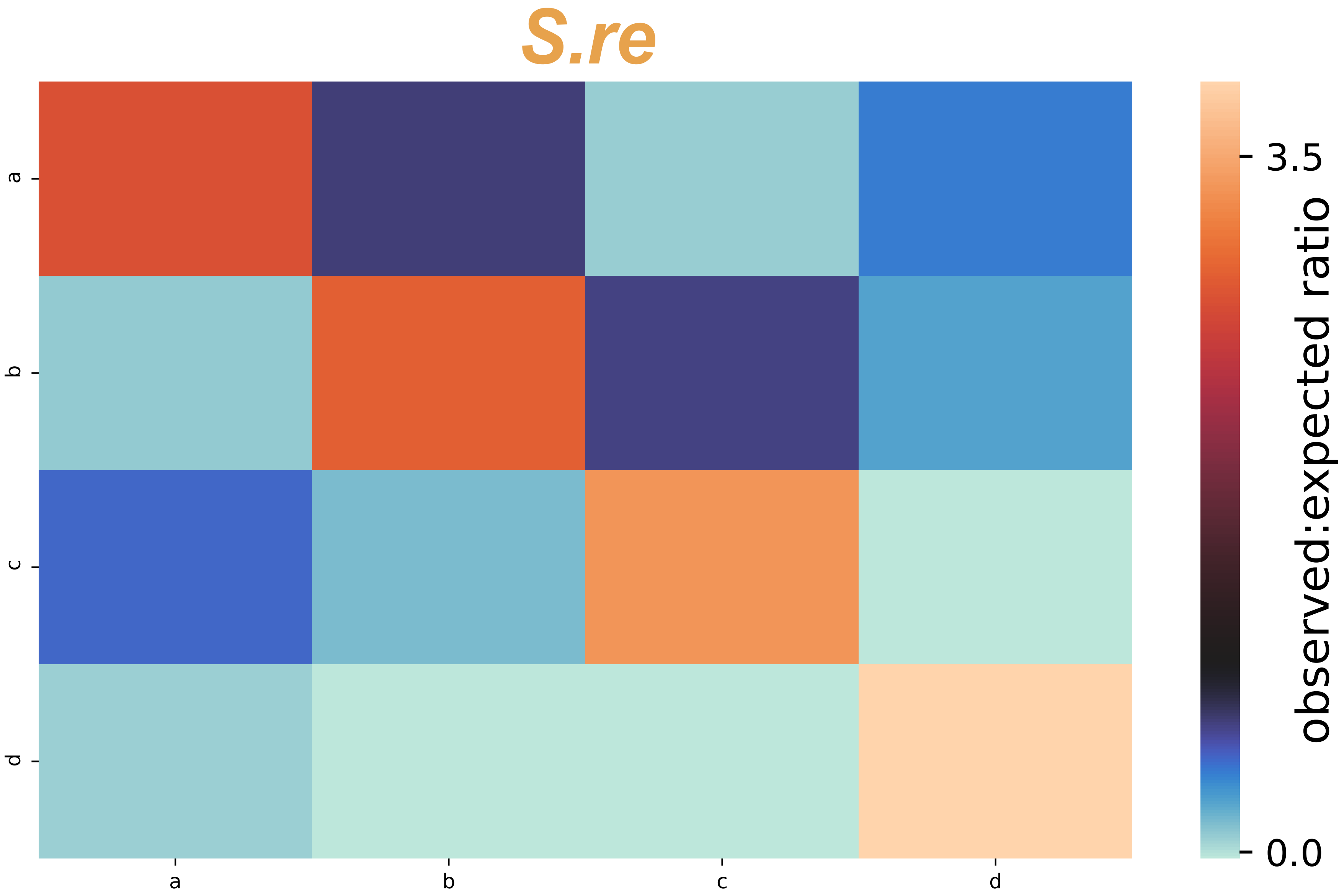

Fig S5

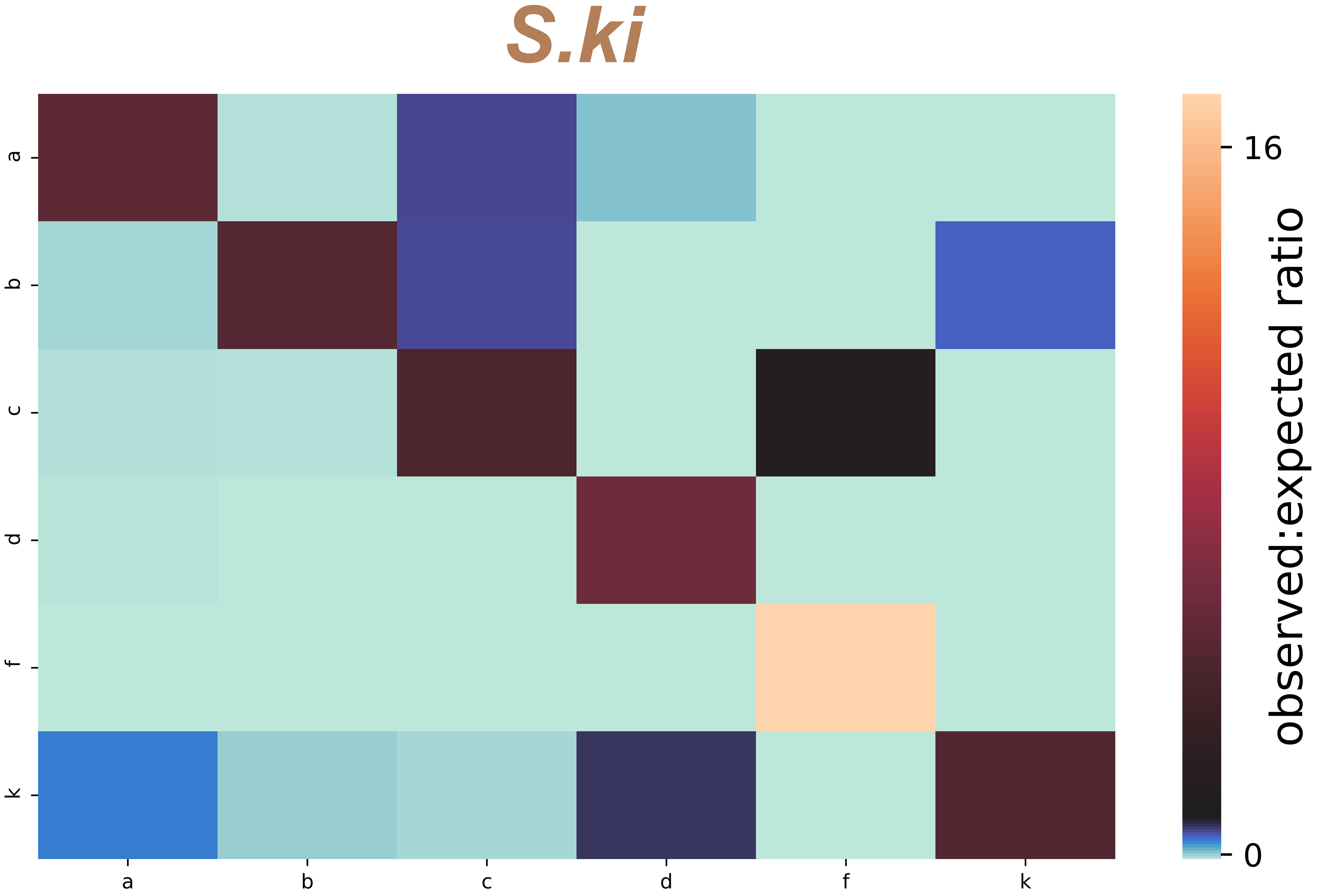

Fig S5

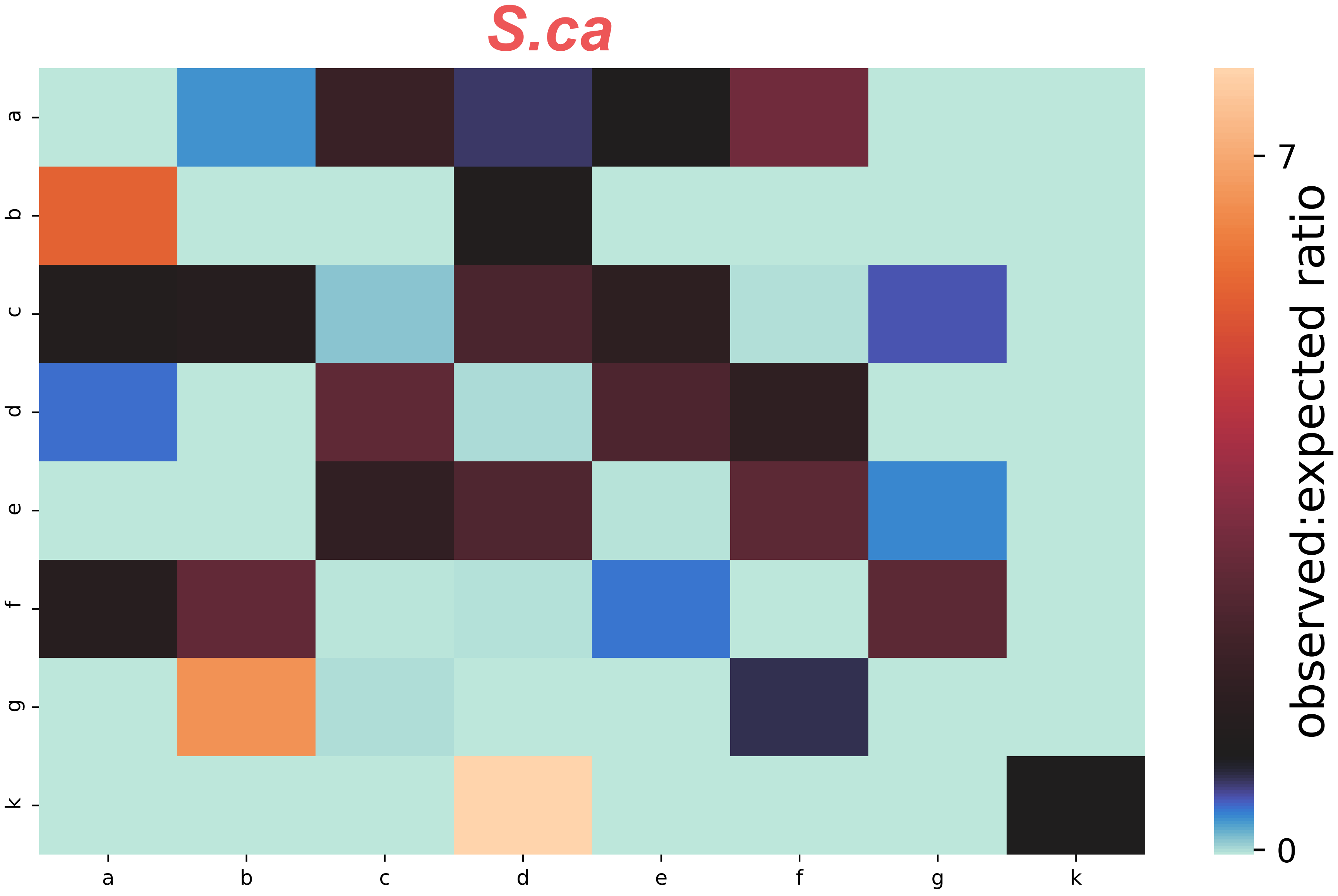

Fig S5

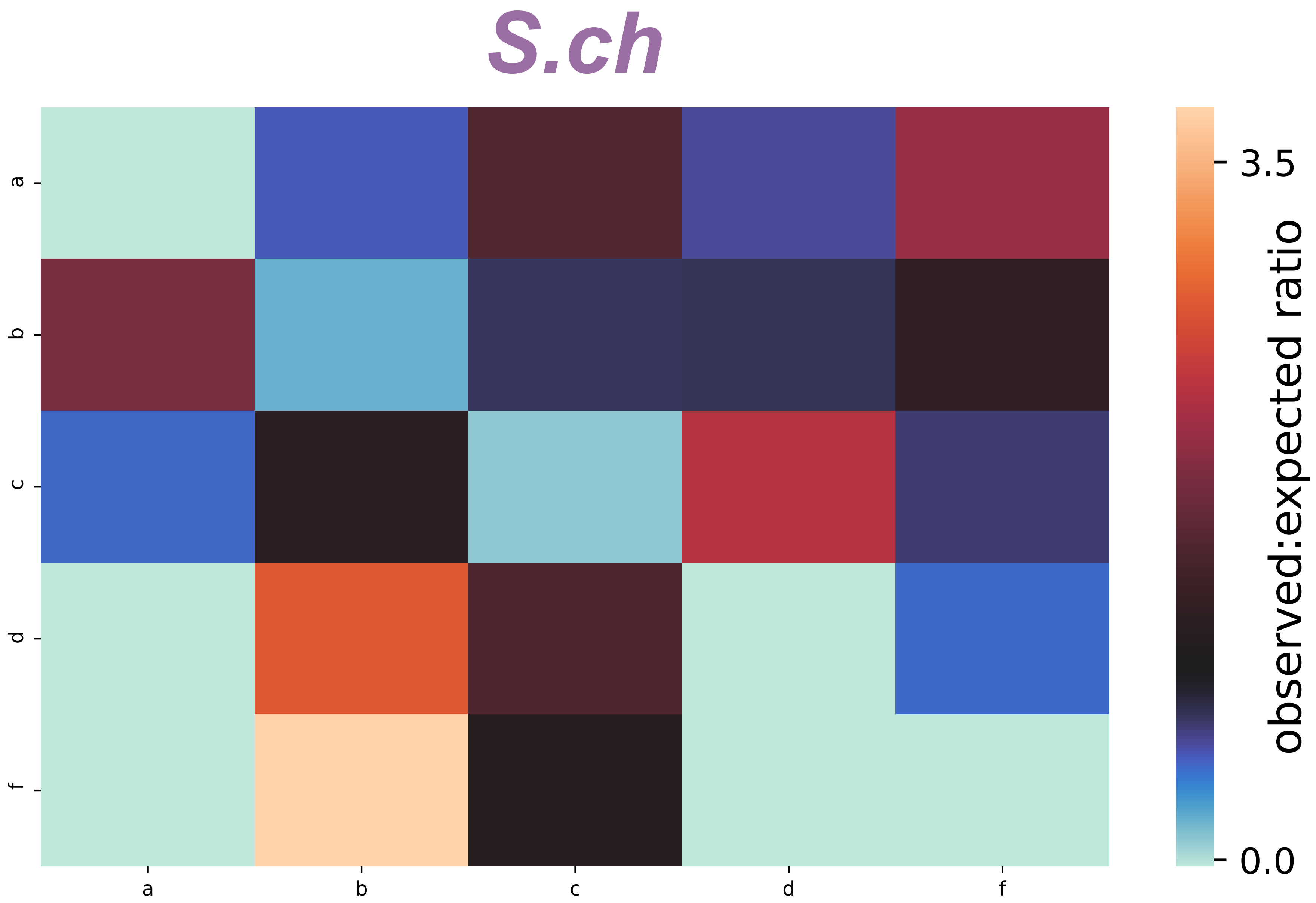

Fig S5

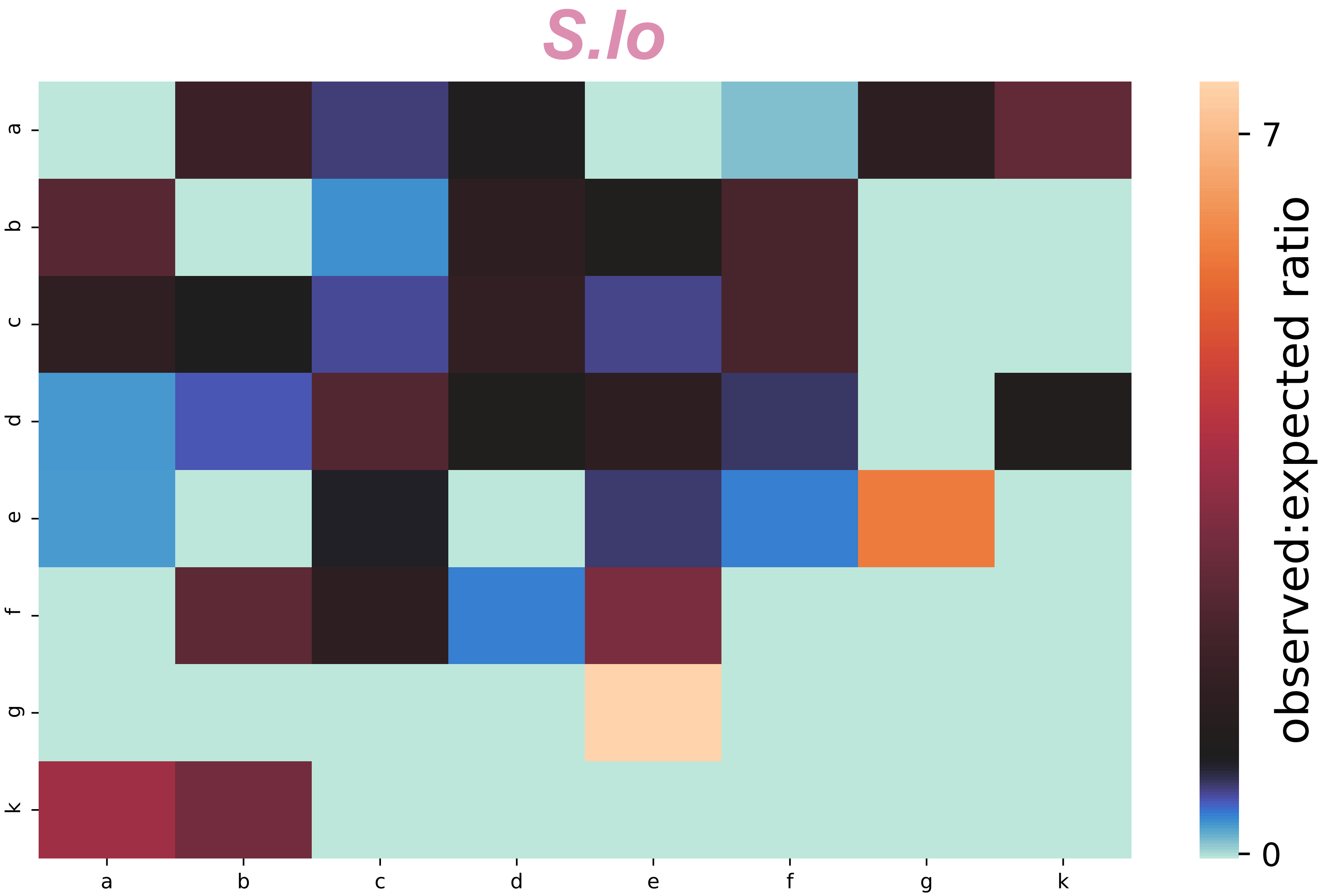

Fig S5

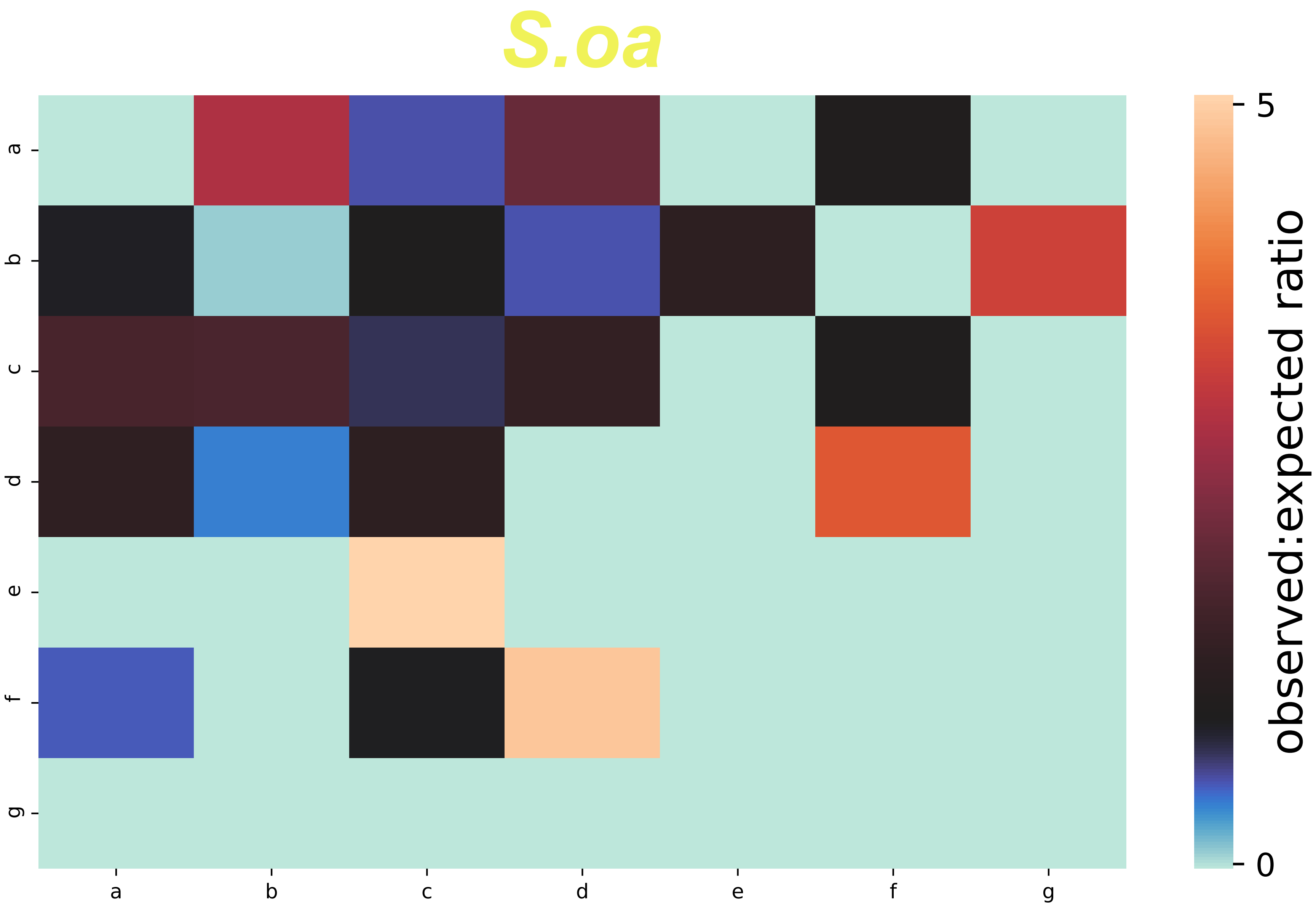

Fig S5

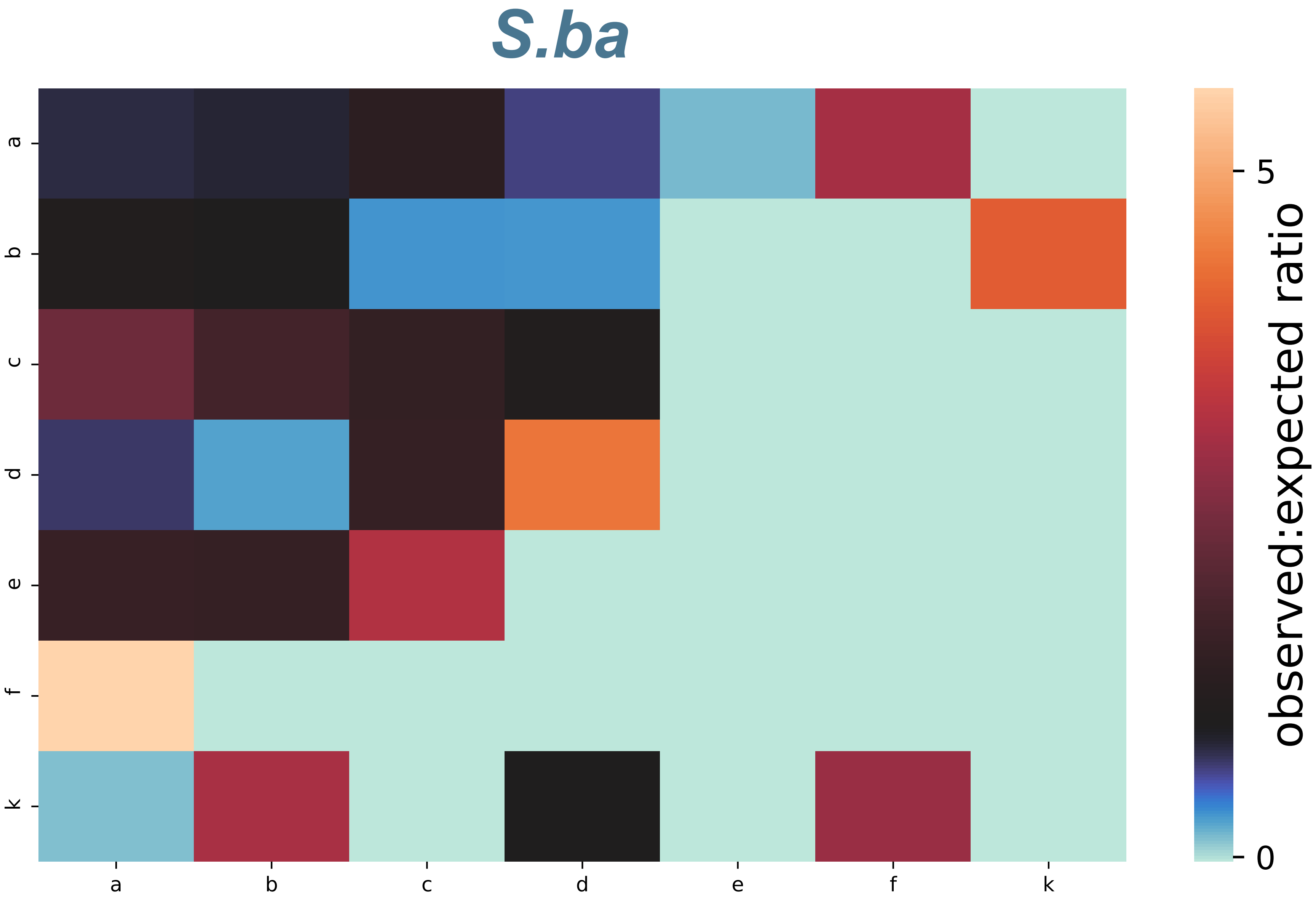

Fig S5

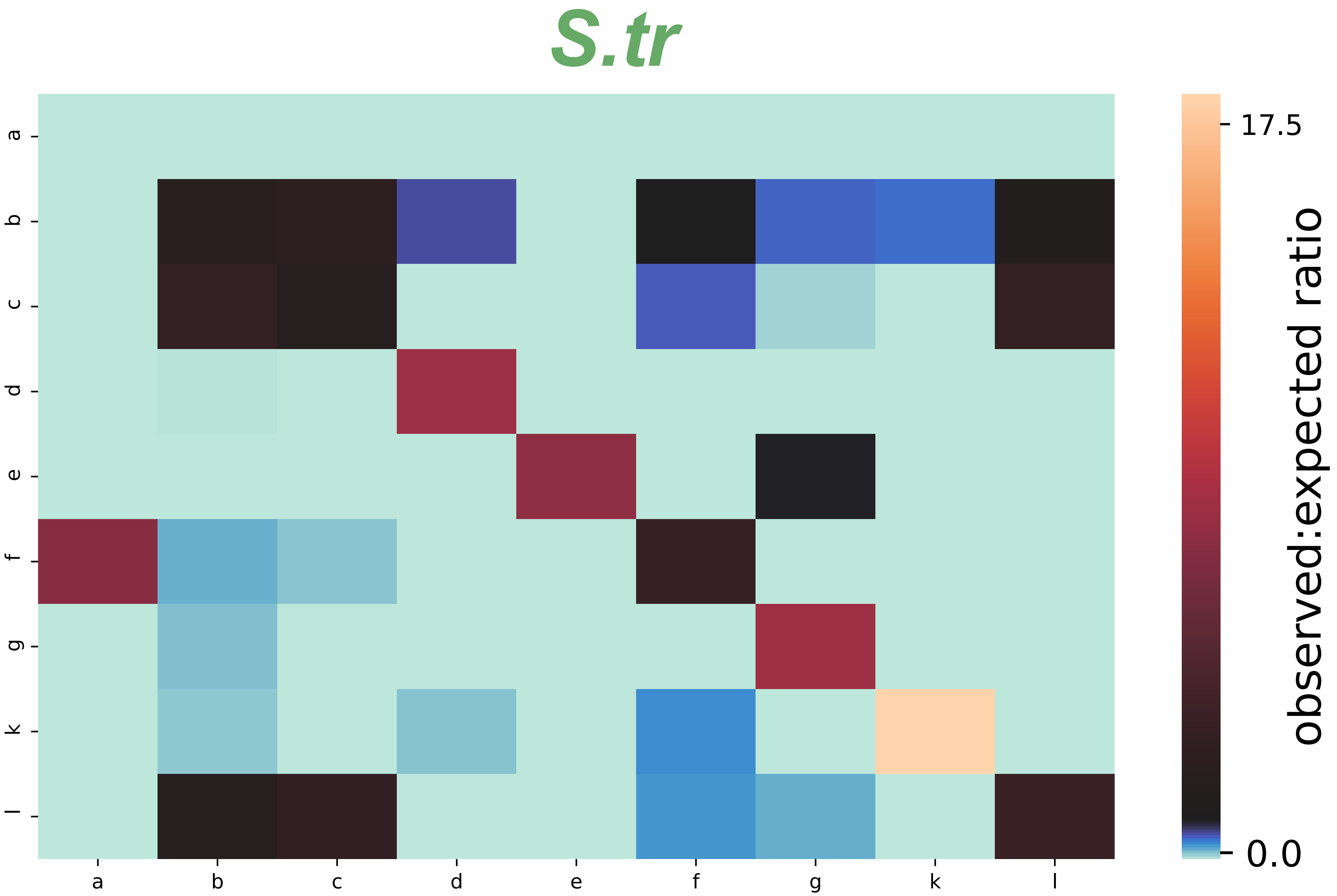

Fig S6

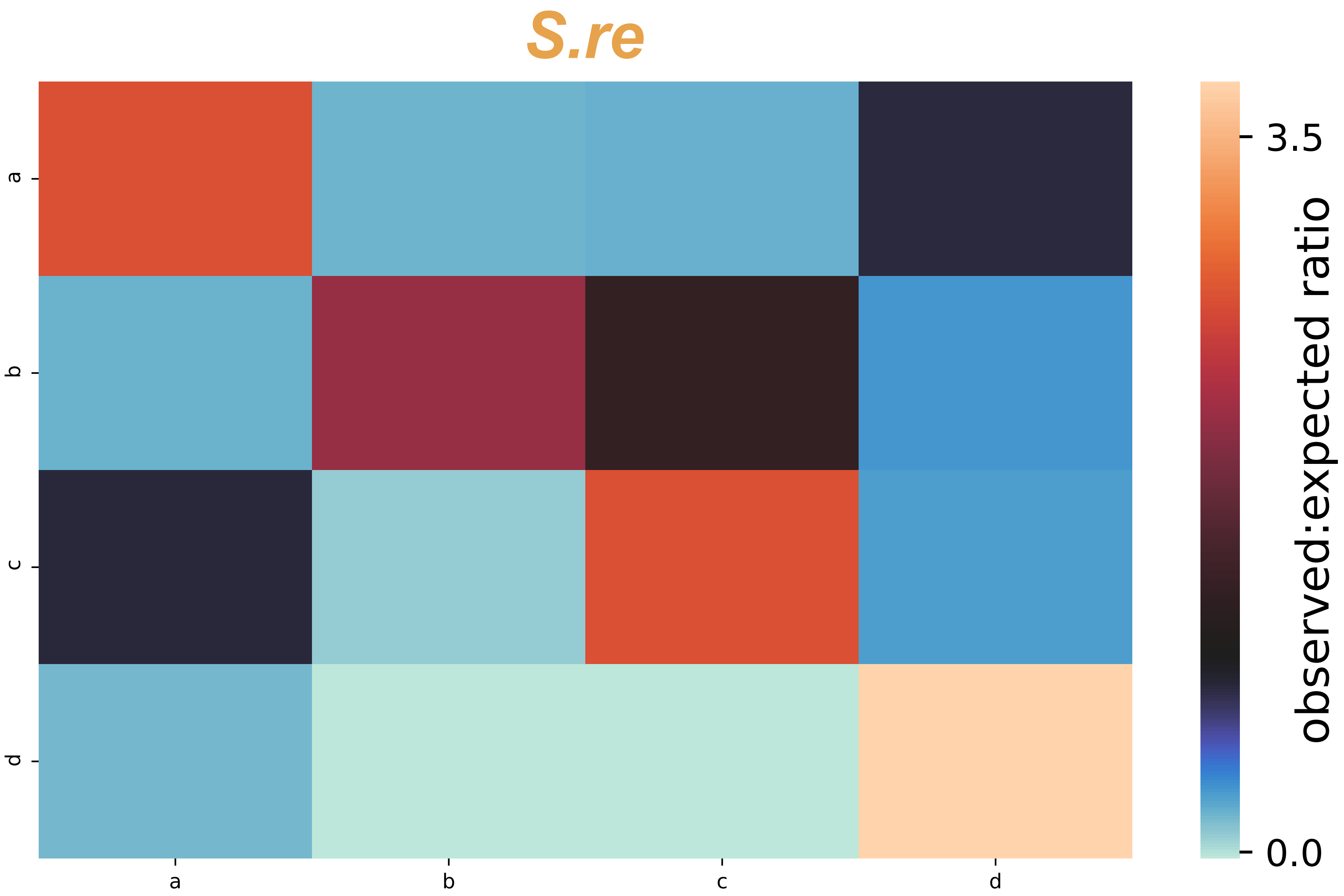

Fig S6

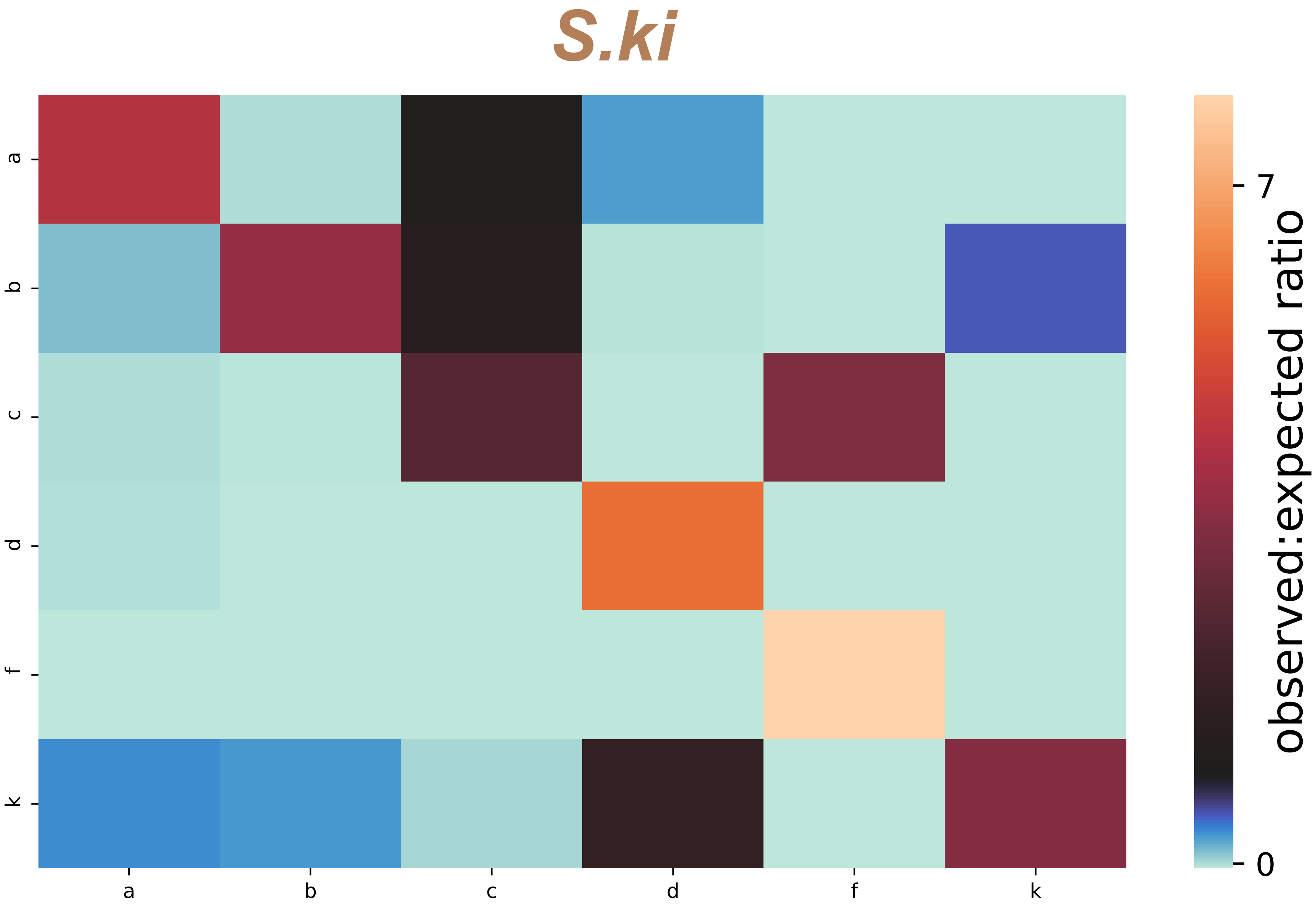

Fig S6

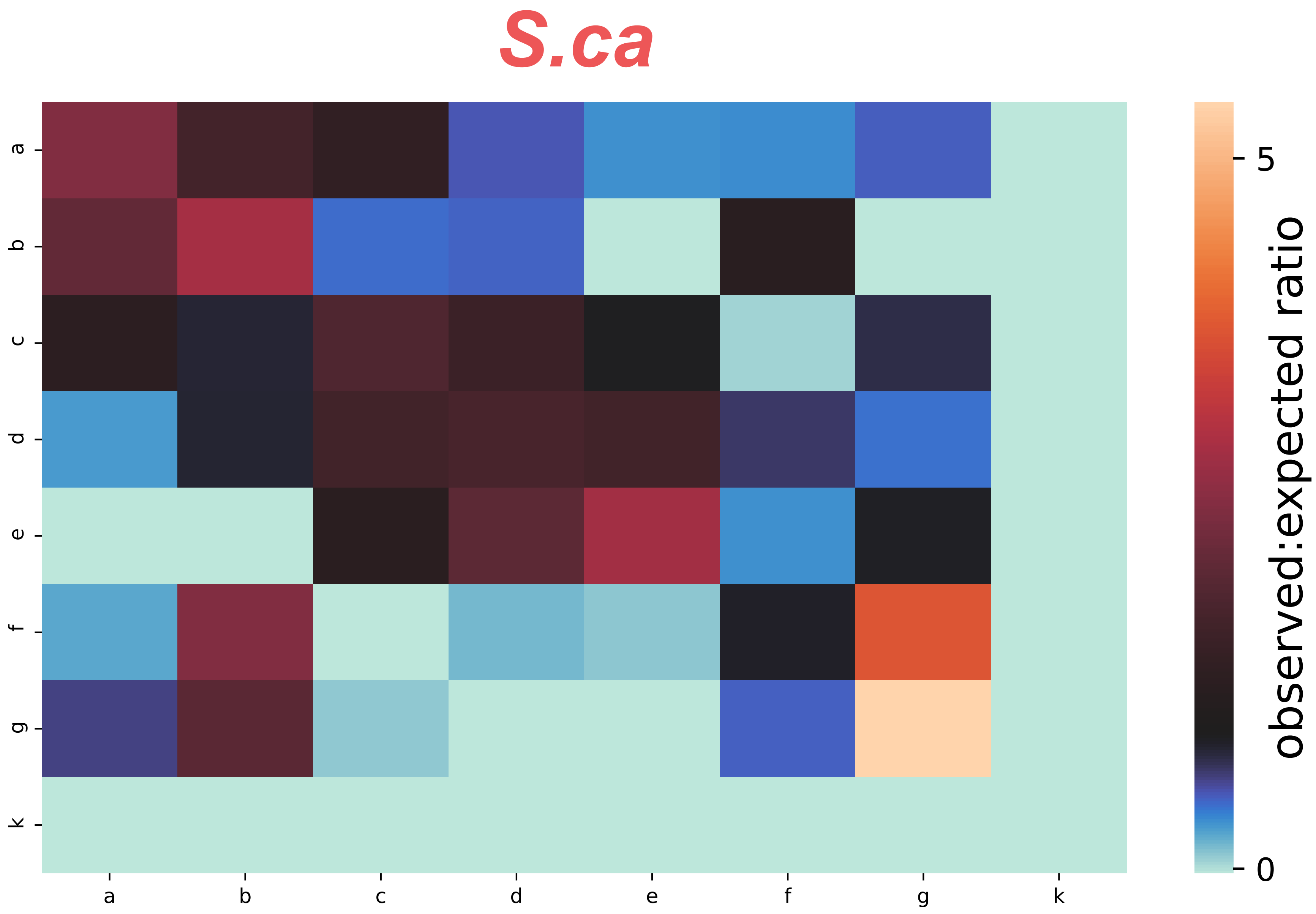

Fig S6

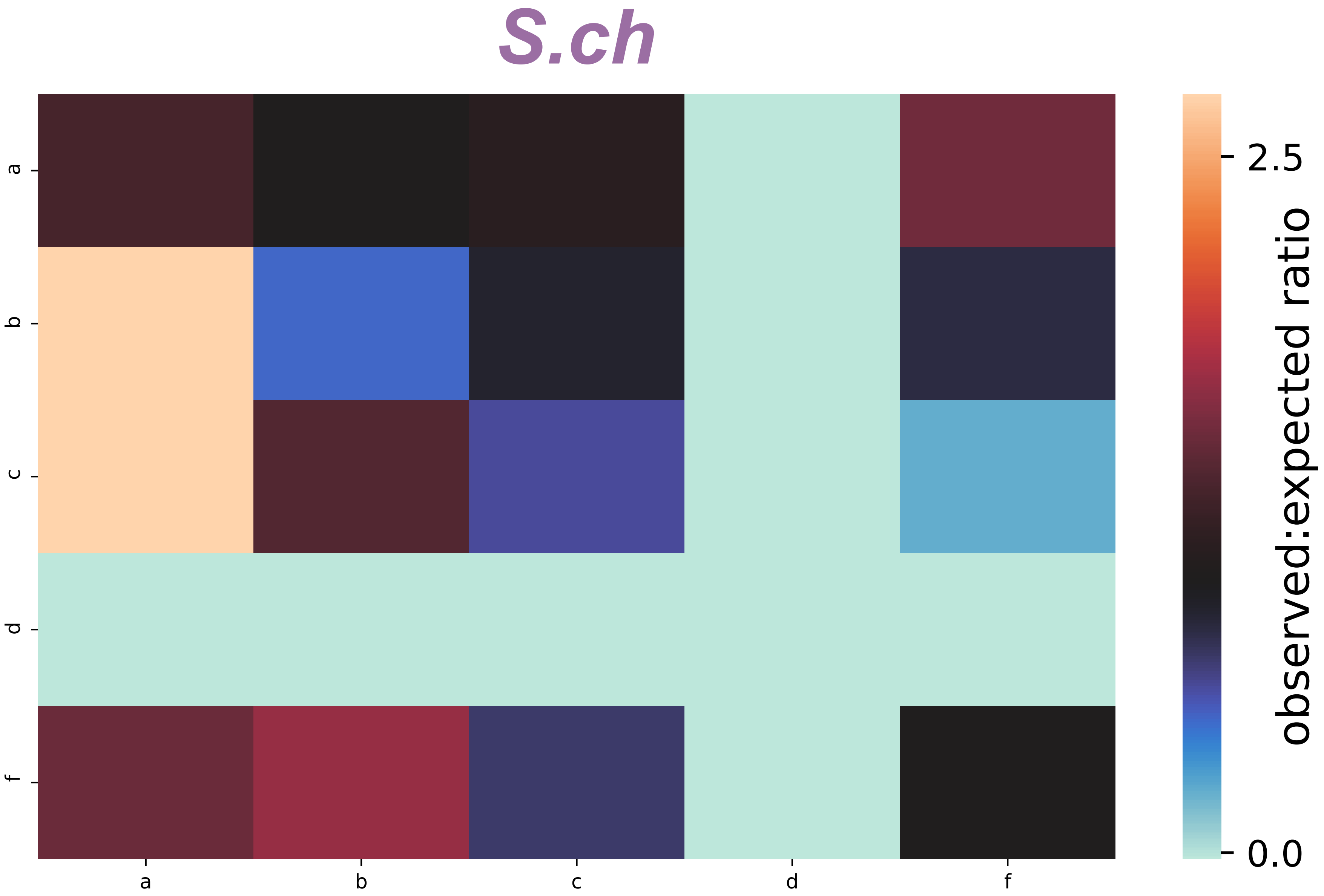

Fig S6

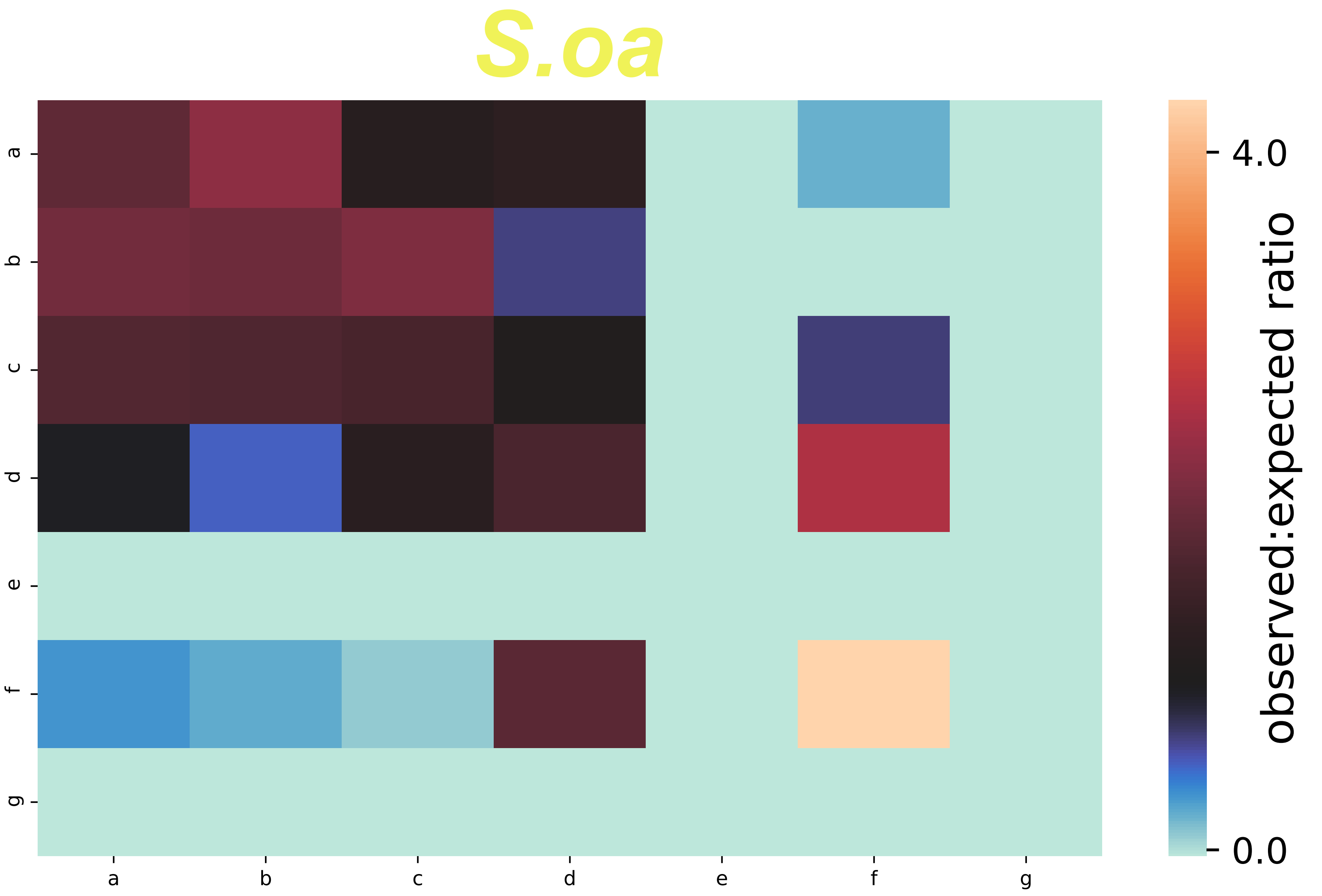

Fig S6

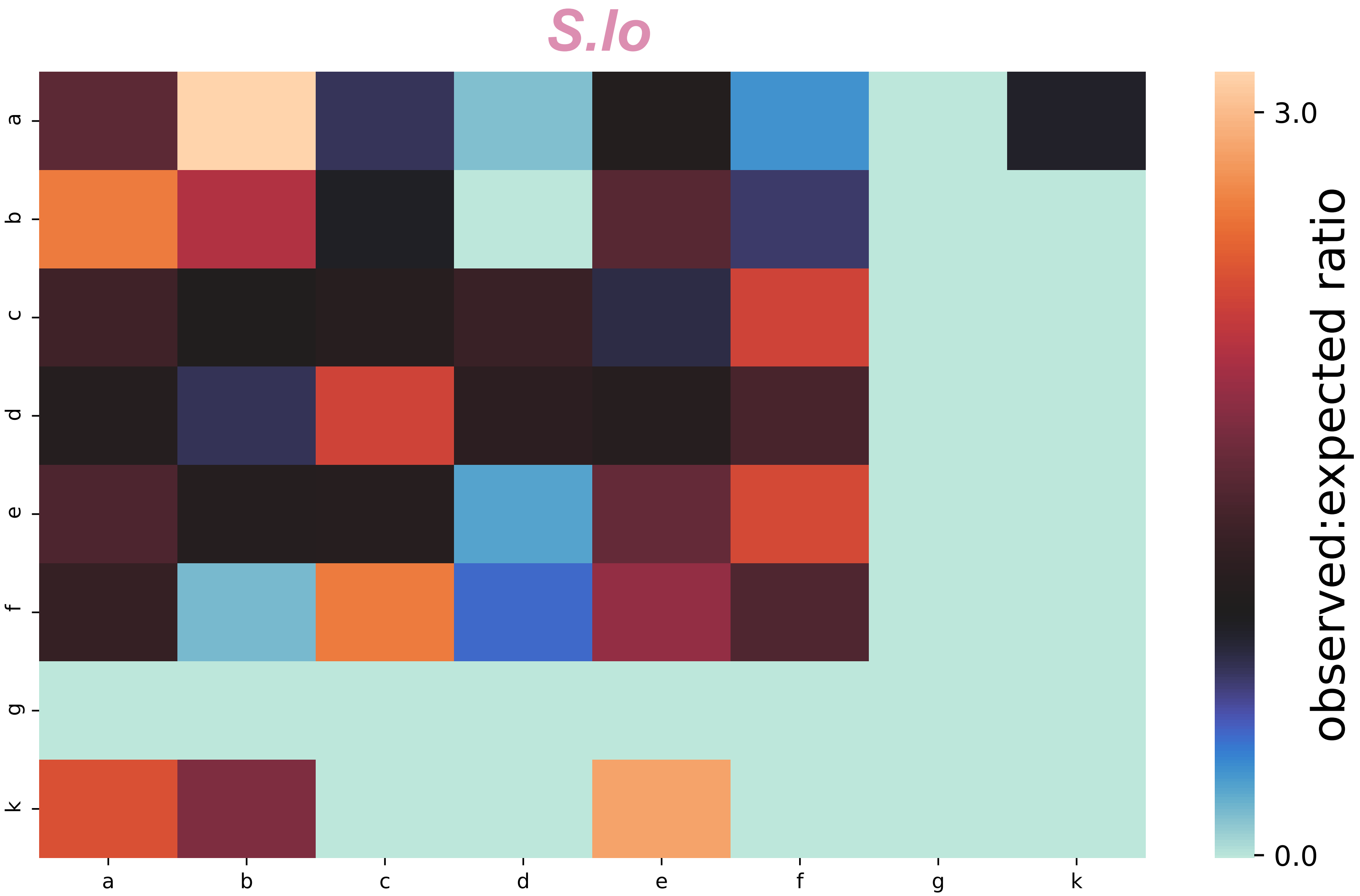

Fig S6

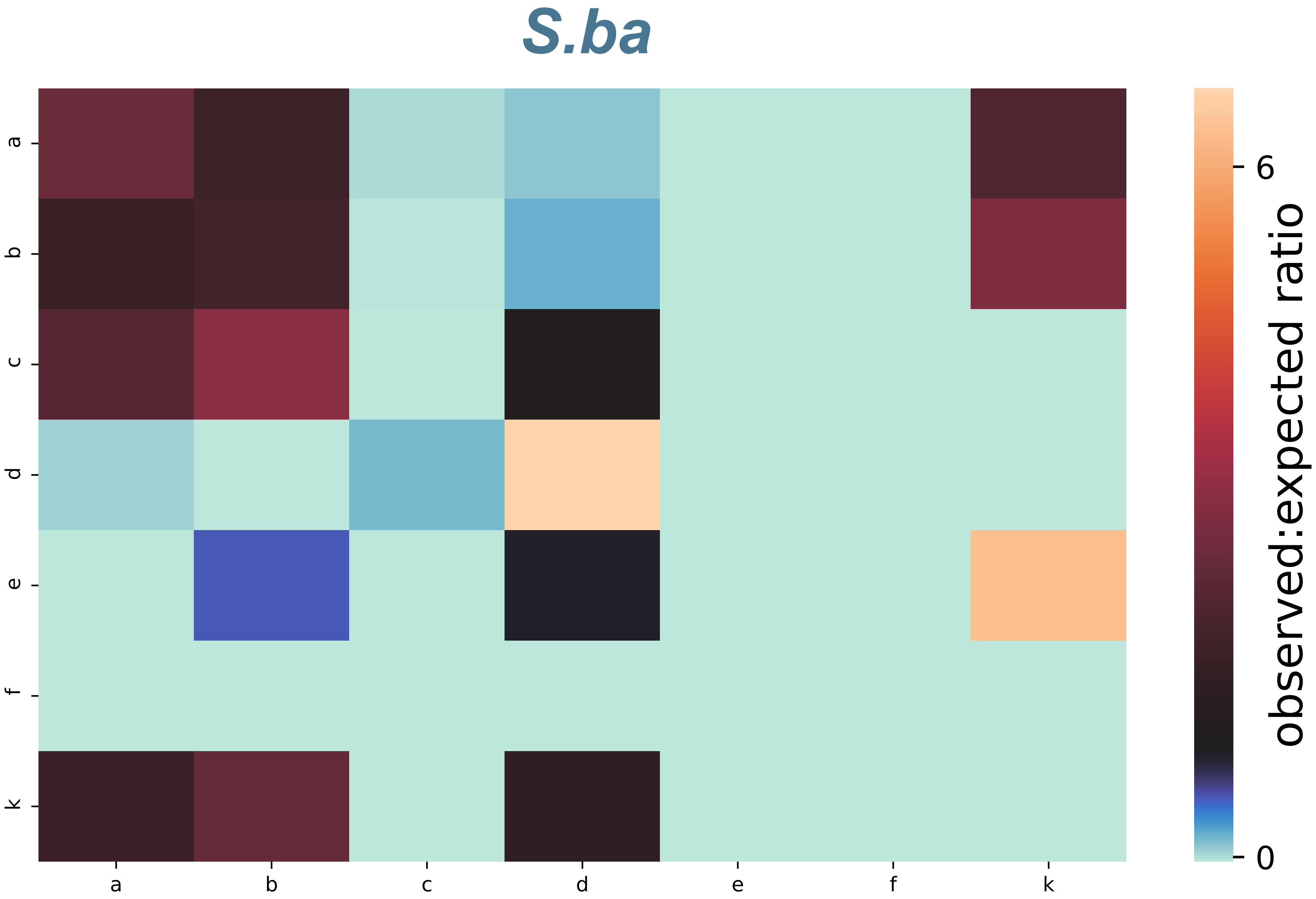

Fig S6

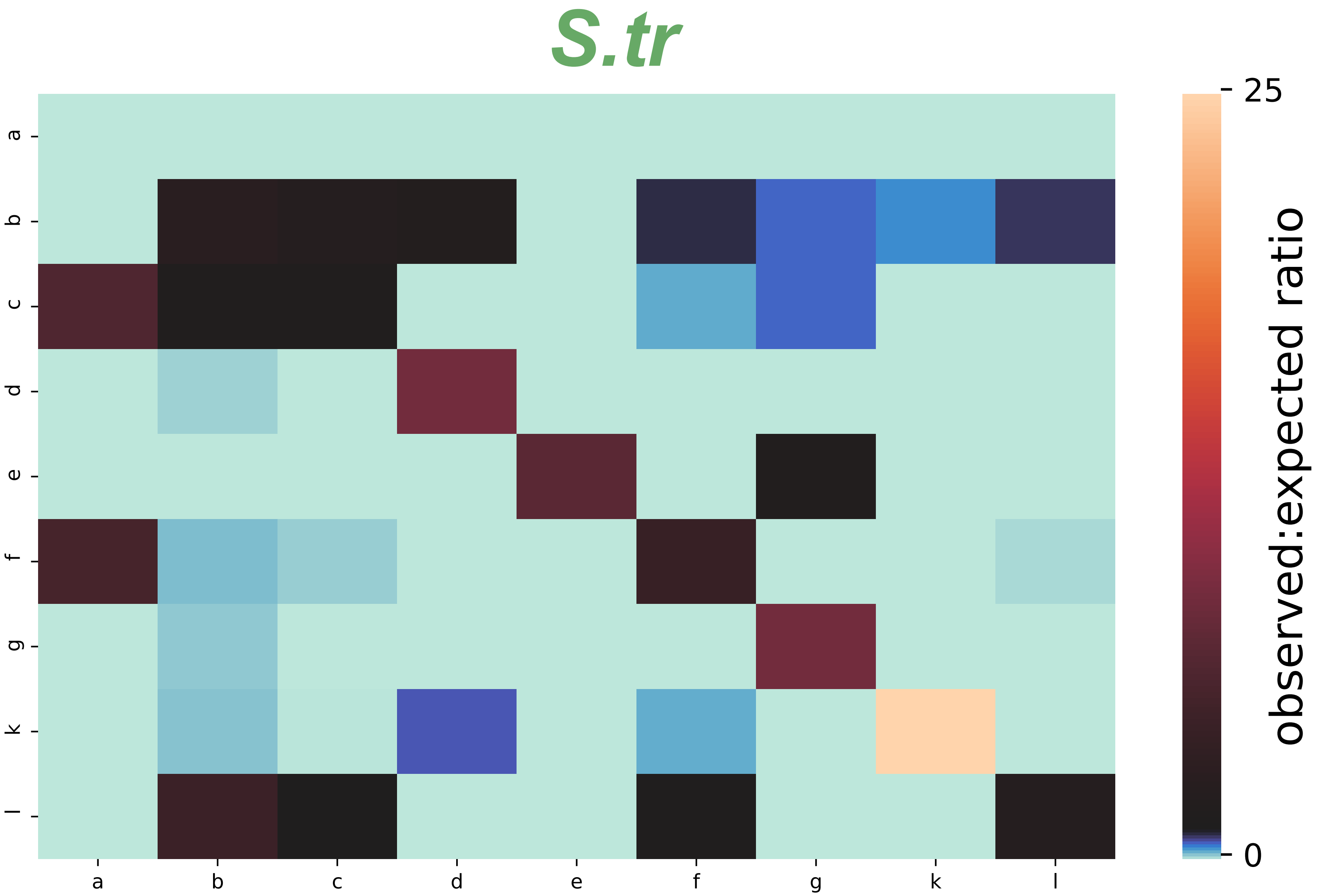

Fig S7

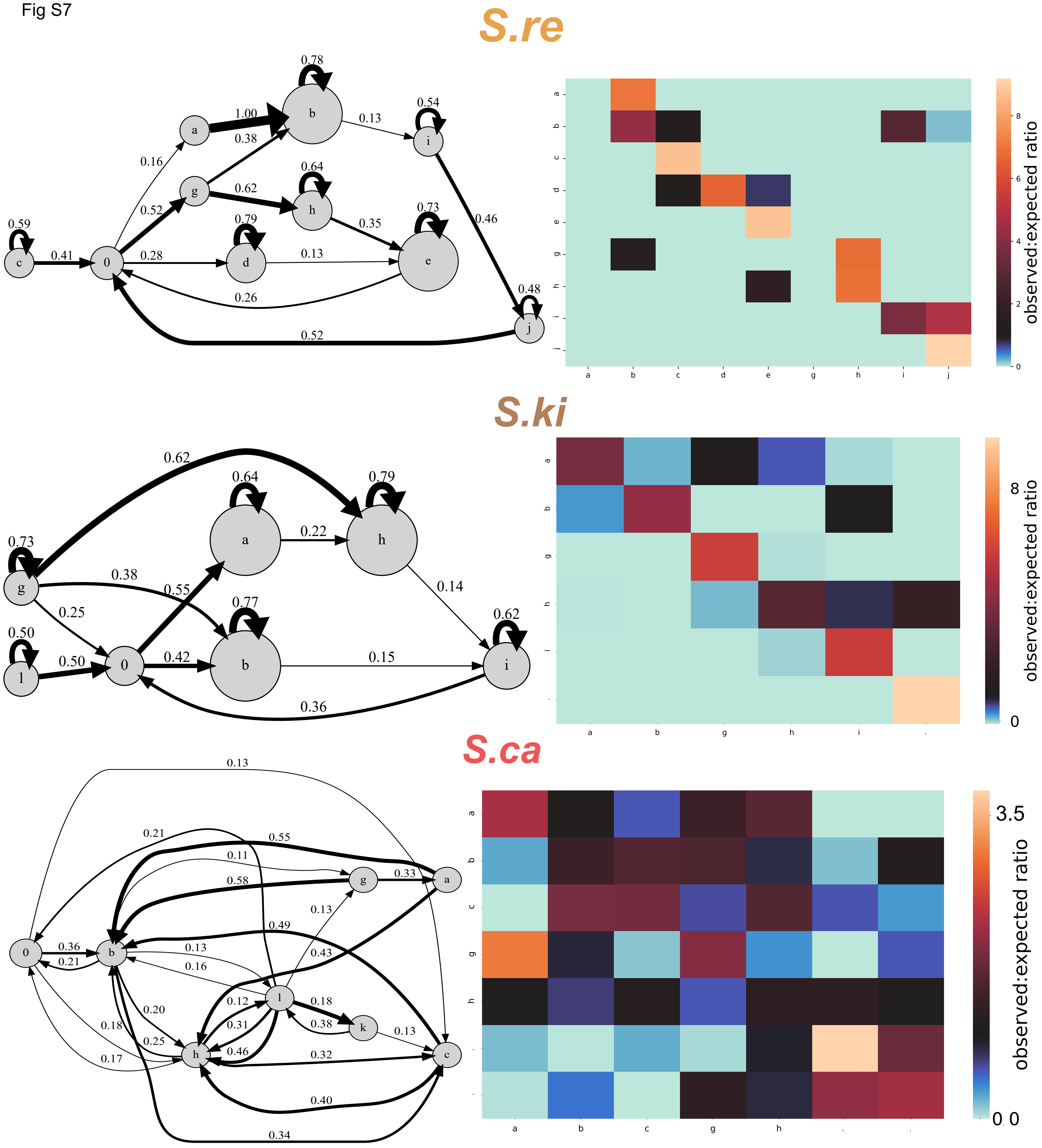
S.re
0.78
1.00
0.38
0.13
0.54
0.16
0.64
0.62
0.52
0.59
0.79
0.35
0.73
0.46
0.41
0.28
0.13
0.26
0.48
0.52
observed:expected ratio
S.ki
0.62
0.64
0.79
0.73
0.22
0.38
0.55
0.14
0.25
0.77
0.62
0.50
0.50
0.42
0.15
0.36
observed:expected ratio
S.ca
0.13
0.21
0.55
0.11
0.58
0.33
0.13
0.49
0.36
0.21
0.13
0.16
0.43
0.20
0.12
0.18
0.18
0.31
0.38
0.13
0.25
0.46
0.17
0.32
0.40
0.34
3.5
0.0
observed:expected ratio

Fig S7

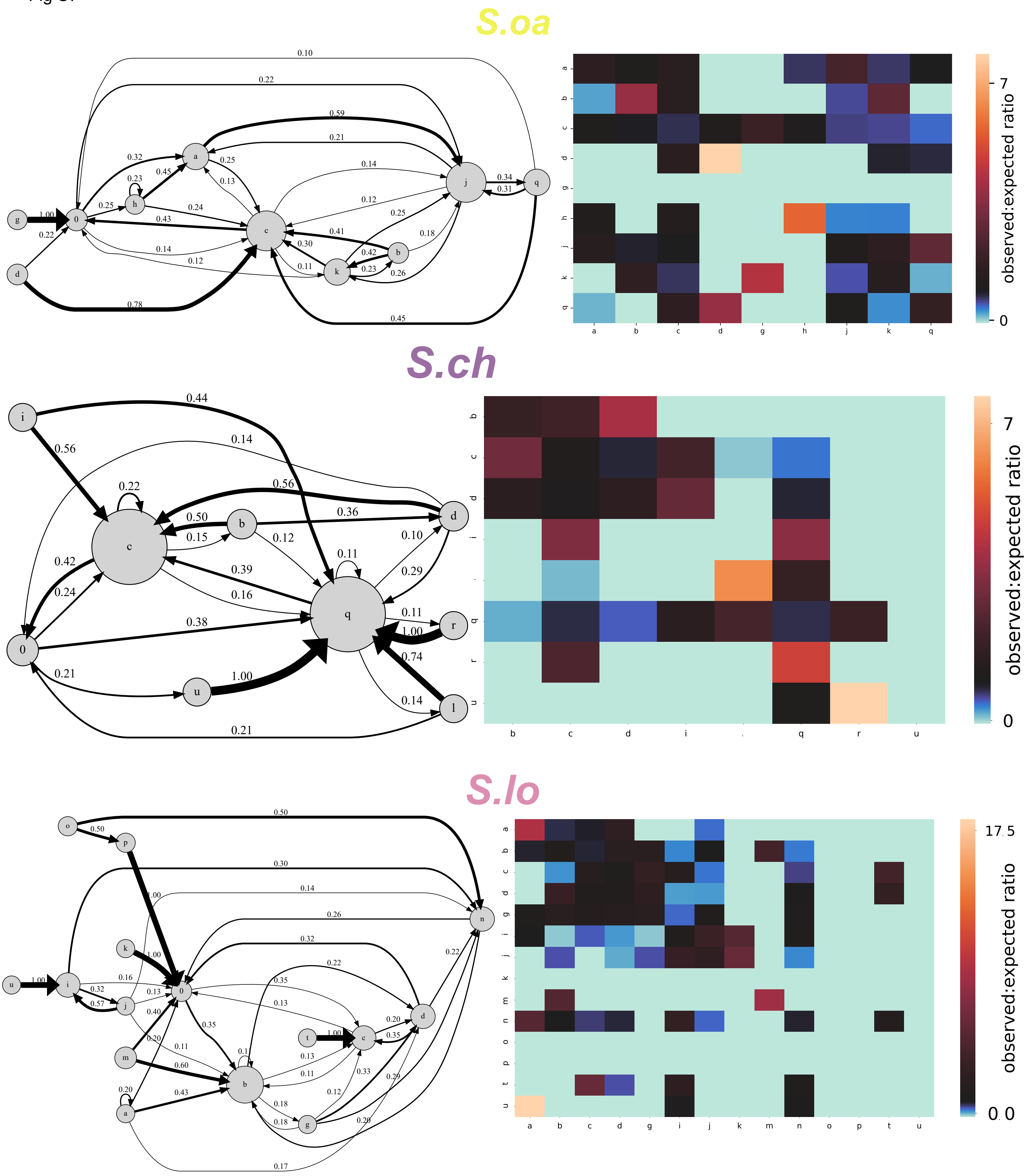

S.oa
observed:expected ratio
S.ch
observed:expected ratio
S.lo
observed:expected ratio

Fig S7

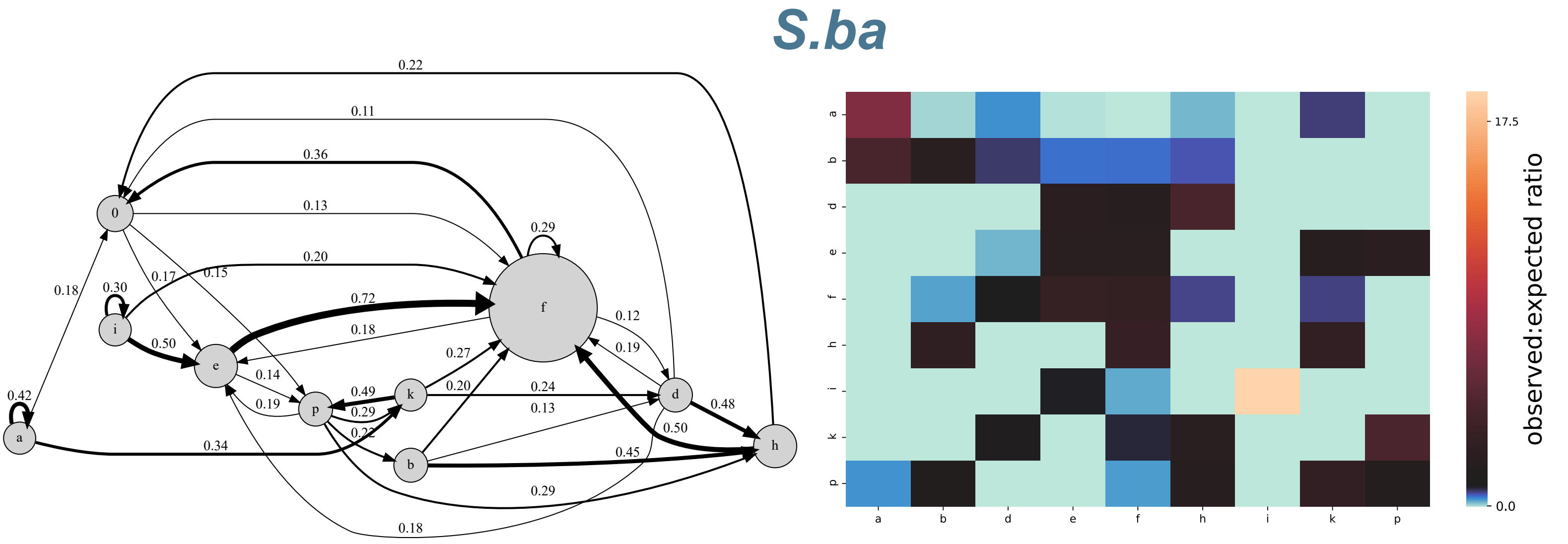
S.ba
observed:expected ratio
17.5
0.0

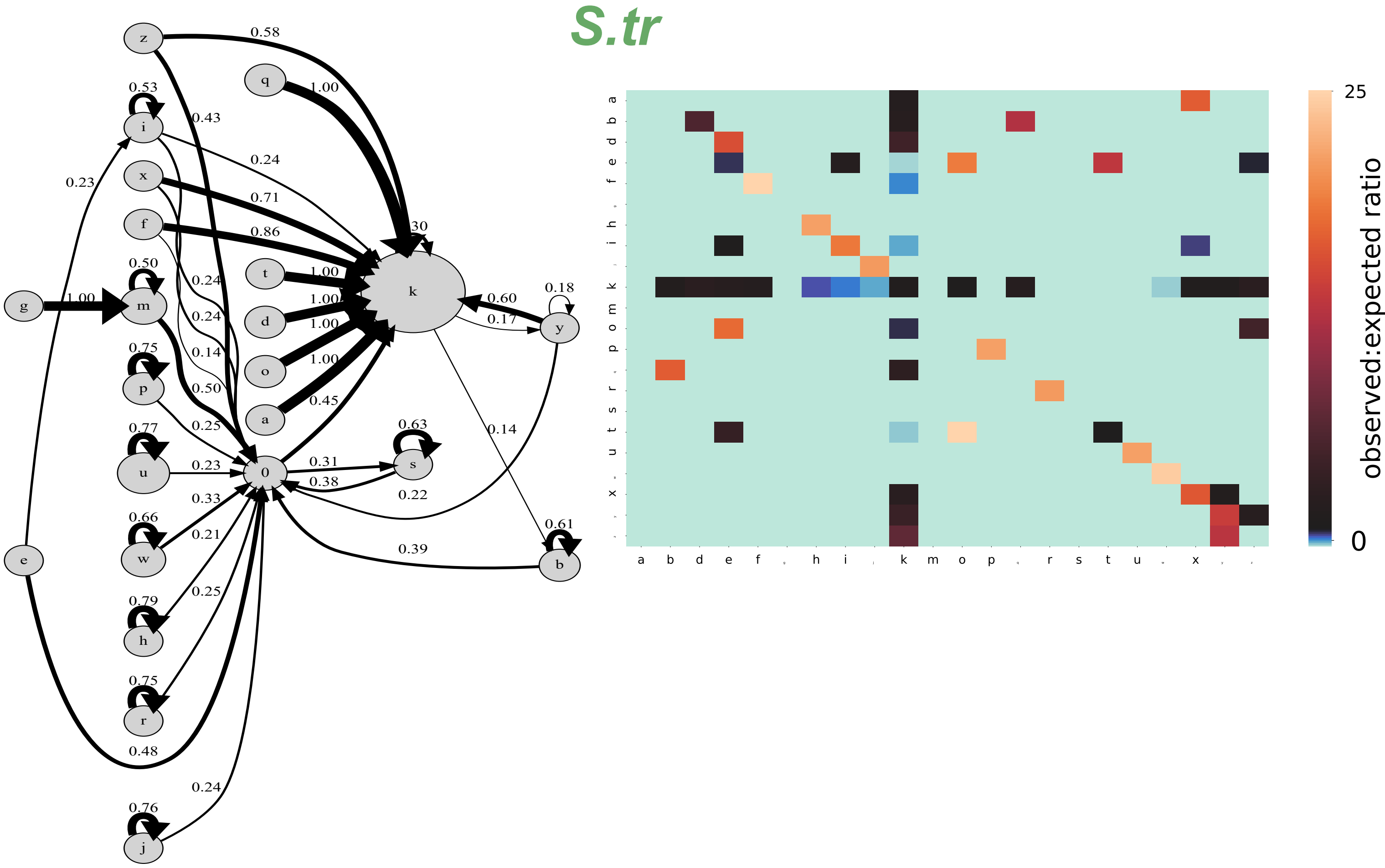
S.tr
observed:expected ratio
25
0

Fig S8

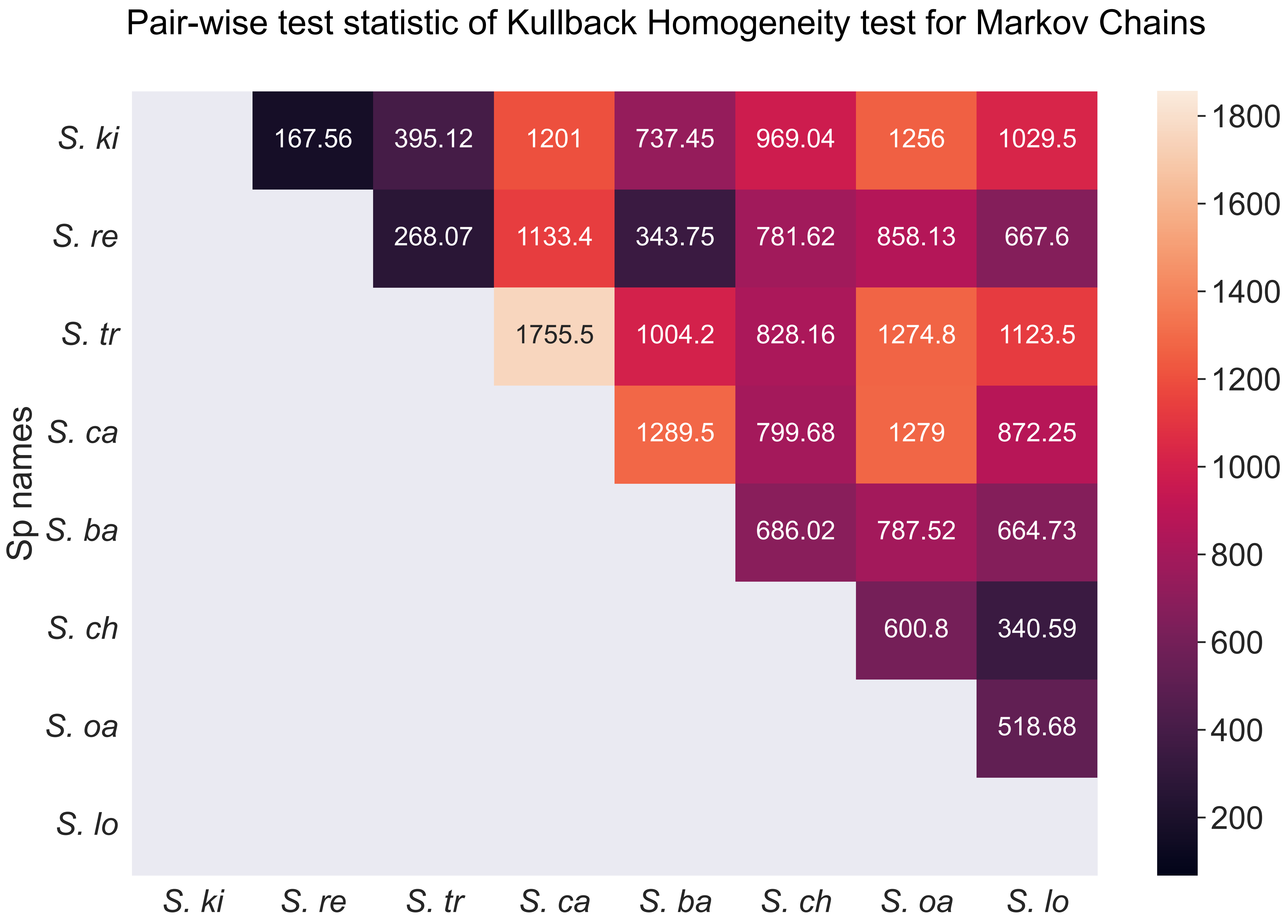


Fig S8: Heat map depicting the pairwise test statistic of the homogeneity test. Species geographically closer to each other generally exhibit a smaller value of the test statistic, suggesting they are more similar to each other than to other congeners.
